# A hippocampus-accumbens code guides goal-directed appetitive behavior

**DOI:** 10.1101/2023.03.09.531869

**Authors:** Oliver Barnstedt, Petra Mocellin, Stefan Remy

**Affiliations:** Department of Cellular Neuroscience, Leibniz Institute for Neurobiology, 39118 Magdeburg, Germany; German Center for Neurodegenerative Diseases (DZNE), 53127 Bonn & 39120 Magdeburg, Germany; International Max Planck Research School for Brain & Behavior (IMPRS), 53175 Bonn, Germany; Center for Behavioral Brain Sciences (CBBS), 39118 Magdeburg, Germany; German Center for Mental Health (DZGP), Magdeburg site, 39106 Magdeburg, Germany

**Keywords:** hippocampus, nucleus accumbens, spatial memory, place cell, reward learning, conjunctive coding, appetitive behavior, calcium imaging

## Abstract

Neurons in dorsal hippocampus (dHPC) encode a rich repertoire of task-relevant environmental features, while downstream regions such as the nucleus accumbens (NAc) translate this information into task-adaptive behaviors. Yet, the contents of this information stream and their acute role in behavior remain largely unknown. Here, we used *in vivo* dual-color two-photon imaging to simultaneously record from both large numbers of hippocampal neurons and an identified subpopulation of NAc-projecting neurons during goal-directed navigation towards a hidden reward zone. We found that NAc-projecting neurons contain enriched spatial information and enhanced representations of non-spatial task-relevant behaviors such as deceleration and appetitive licking, both of which could be elicited by optogenetic activation of dHPC terminals in NAc. A generalized linear model revealed enhanced conjunctive coding in NAc-projecting dHPC neurons, improving the identification of the reward zone. We propose that dHPC routes specific reward-related spatial and behavioral state information to guide NAc action selection.

## Introduction

Memories allow an organism to use past experience to optimize current and future behaviors (1). The hippocampus (HPC) is widely recognized as one of the main sites of memory-related plasticity (2–4). Yet, while our understanding of memory processing within the HPC has greatly advanced, surprisingly little is known about the translation of this information into behavioral action: which hippocampal output pathways send which types of information to guide memory-driven behavior? This lack of knowledge corresponds with the scarcity of data available on the activity profile of large numbers of identified hippocampal projection neurons while animals perform memory-dependent behaviors.

The HPC is a major part of the brain’s limbic system, processing various kinds of memory by receiving highly processed sensory information from the entorhinal cortex, and sending behaviorally relevant outputs to diverse brain regions via hippocampal Cornu Ammonis field 1 (CA1) and subiculum (5). Its elevated role for spatial memory is highlighted by the presence of spatially tuned “place cells” that form a “cognitive map” to support goal-directed navigation (6, 7). This cognitive map is further supported by neuronal coding of navigationally relevant features such as borders, speed, reward/goal locations (8–11), but also non-spatial information such as that about future decisions or behavioral tasks (12, 13). The conjunction of this variety of encoded features is suggested to provide a “scaffold” that supports the formation of episodic memories (14).

Previous models of hippocampal memory processing assumed largely homogeneous cell populations (15, 16). However, hippocampal principal neurons are increasingly recognized as structurally and functionally diverse in terms of their morphology, electrophysiology, transcriptomic cell types, anatomical differences across its three axes, and projection patterns (17, 18). Such heterogeneity may provide the hippocampus with the intrinsic flexibility to meet the various demands of diverse environments (17), and to route task-relevant information to specific output targets (19, 20).

One target receiving strong projections from both dorsal and ventral CA1 and subiculum is the nucleus accumbens (NAc) (21, 22), a basal ganglia brain structure crucial for value-based action selection (23, 24). Described as the “interface between limbic and motor circuitry” (25), it has been suggested as the main site of transformation from a hippocampal spatial code into a motivation-driven motor code (26). In line with this, NAc neurons are spatially tuned, required for spatial memory acquisition and consolidation, and display task-dependent synchrony with dHPC neurons (27–30). Indeed, disabling HPC → NAc projections diminished conditioned place preference (CPP) (31, 32), and optogenetic stimulation was sufficient to artificially induce CPP (33, 34). While these findings cement the NAc’s role as an indispensable hippocampus output node, surprisingly little is known about which spatial, contextual and behavioral patterns of hippocampal information the NAc receives. Here, we set out to understand the specific contents of this information stream during goal-directed navigation and its acute role in behavior. We hypothesized that the HPC may selectively route both spatial as well as other task-relevant information to aid the NAc in action selection.

To test this, we employed dual-color two-photon imaging, capturing *in vivo* calcium signals of large populations of neurons in dHPC, while using a projection-specific red fluorophore to allow identification of NAc-projecting neurons (dHPC^→NAc^). Directly comparing dHPC^→NAc^ activity with the rest of the dHPC population (dHPC^−^), we found enhanced spatial tuning in dHPC^→NAc^ neurons, with strong modulation by local cue boundaries and a reward zone. We also found elevated coding of low velocities and appetitive licking, both of which could be elicited by optogenetic stimulation. Lastly, we show stronger conjunctive coding of space, velocity, and appetitive licking, which improves classification of the reward zone with a linear decoder. We thus propose that dHPC routes reward context-enriched spatial and behavioral state information to bias NAc action selection.

## Results

### Dual-color calcium imaging during goal-directed navigation

To investigate coding of spatial information and goal-directed behavior in dHPC^→NAc^ neurons, we trained food-restricted mice on a head-fixed spatial reward learning task. Mice had to run on a self-propelled treadmill, traversing a 360 cm long textile belt with six differently textured zones, including one otherwise unmarked 30 cm long fixed reward zone (35, 36). Licking a spout in this zone causes a liquid reward to be dispensed once per lap (Figure 1A and Figure S1A-E). Mice underwent five days of training in which they learned to obtain more rewards by progressively increasing their licking in both reward zone and a 30 cm “anticipation zone” preceding the reward zone (Figure 1B-D and Figure S1F-N).

**Figure 1.**
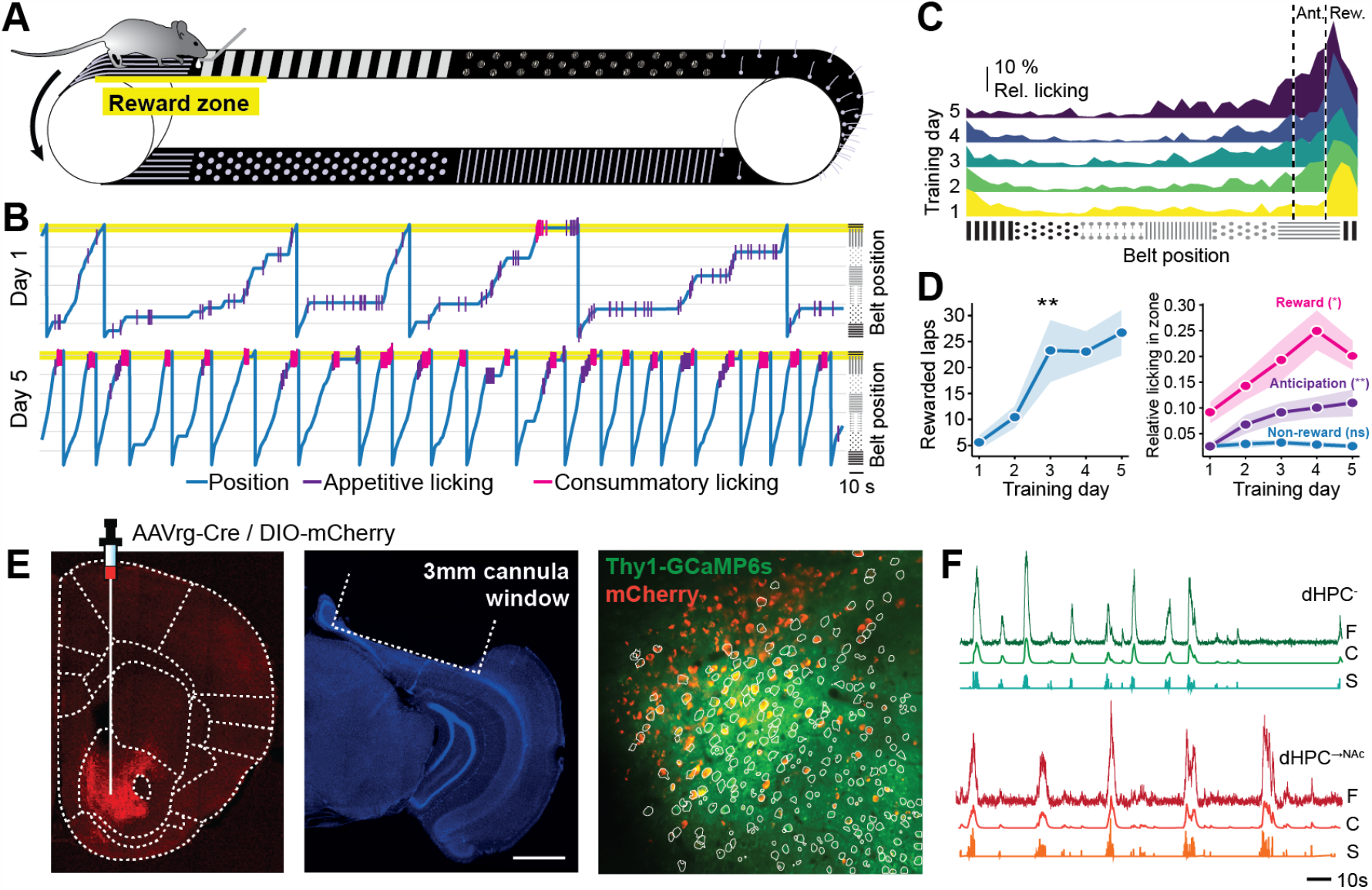
Dual-color two-photon calcium imaging of dHPC allows projection-specific activity monitoring during goal-directed navigation. (A-D) Training mice on a spatial reward learning task. (A) Schematic of behavioral task. Mice moved on a cue-enriched self-propelled treadmill belt of 360 cm length and obtained a liquid reward when licking at a spout in a hidden reward zone of 30 cm length (yellow). (B) Representative traces of one mouse’s behavior on training day 1 and day 5. (C) Licking in anticipation (Ant.) and reward (Rew.) zone increases over the course of training. (D) Number of rewarded laps per training session (*F* (4) = 6.344, GG-correction), anticipatory (*F* (4) = 3.803) and reward licking (*F* (4) = 4.276, GG-correction) significantly increased over the course of five training days (*n* = 18 mice, repeated-measures ANOVA). See also Figure S1. (E) Schematic of dual-color projection neuron imaging method. Thy1-GCaMP6s mice were injected with AAVrg-Cre in the medial NAc and DIO-mCherry in dHPC. Representative coronal brain slice showing axonal mCherry expression in NAc (AP -1.3; left). Representative coronal brain slice stained with DAPI (blue) of dHPC, showing the outlines of the 3 mm cannula window used for imaging; scale bar represents 1 mm (second left). Field of view of one sample experiment showing Thy1-GCaMP6s expression in green and mCherry expression of putative NAc-projecting neurons in red; outlines show detected components used for analysis (right). (F) Two representative neurons’ raw (“F”), denoised and deconvolved (“C”) and event (“S”) traces; red traces indicate mCherry co-expression (dHPC^→NAc^). See also Figure S2. All data are presented as mean ± SEM. \**p* < 0.05, \*\**p* < 0.01.

To understand the neural coding properties of large numbers of dHPC^→NAc^ neurons in behaving animals, we turned to dual-color two-photon imaging in mice pan-neuronally expressing the calcium indicator GCaMP6s, as well as the static red marker mCherry in defined NAc-projecting neurons. For this, we injected AAVrg-Cre in NAc and DIO-mCherry in dHPC (CA1/subiculum border region) of Thy1-GCaMP6s mice (37). This approach allowed us to obtain dynamic calcium signals both in a large majority of mCherry-negative hippocampal neurons (dHPC^−^) and specifically mCherry co-expressing NAc-projecting neurons (dHPC^→NAc^), simultaneously within the same field of view using the same calcium indicator (Figure 1E and Figure S2). It allowed us to overcome constraints of electrophysiological studies such as relatively low sample sizes (19, 20) or indirect connectivity measurements (38, 39). Optical access to dorsal CA1 and pro-subiculum (also known as proximal subiculum (40, 41)) was established by implanting a chronic hippocampal window after virus injections (Figure S1C) (42, 43). Imaging data was acquired after 5 days of behavioral training, was motion-corrected using NormCorre, and spatio-temporal components were extracted using constrained non-negative matrix factorization (CNMF) (44). We thus obtained calcium signals for further analysis from a total of 5,372 GCaMP-expressing neurons including 444 putative dHPC^→NAc^ neurons in 6 mice across 19 imaging sessions (Figure 1F and Figures S2 and S3). These numbers approximate previously established proportions of NAc-projecting neurons in dorsal prosubiculum and distal CA1 (20, 45, 46).

### Enhanced spatial coding by dHPC^→NAc^ neurons

Given the dHPC’s well-established role in representing spatial information and the role of dHPC^→NAc^ projections in spatial memory tasks (31, 32), we wondered if and how spatial information may be encoded by dHPC^→NAc^ neurons compared to dHPC^−^ neurons. For this, we extracted calcium events for each neuron and compared this activity across the linear spatial environment of the treadmill belt as mice traversed it lap by lap. Indeed, we found large numbers of both dHPC^−^ and dHPC^→NAc^ neurons with repeatedly elevated calcium levels on the same positions of the belt (Figure 2A-C and Figure S3). Comparing each neuron’s spatial information content (47) with that of a randomly shuffled distribution, we identified a significantly higher proportion of such place cells in dHPC^→NAc^ neurons (169/444 neurons; 38 %) compared to dHPC^−^ neurons (1,581/4,928 neurons; 32 %; χ^2^, *p* = 0.012; Figure 2D). Within this population of place cells, we further analyzed how specifically space is encoded, and found that dHPC^→NAc^ place cells had a higher spatial information rate (47) (*p* < 0.001, Welch’s *t*-test; Figure 2E). They also had significantly higher levels of sparsity (*p* < 0.001, Welch’s *t*-test; Figure 2F), a measure for how diffuse a neuron is firing in the spatial domain (48). Furthermore, both the relative calcium activity (activity inside the place field – activity outside the place field; *p* = 0.0044, Welch’s *t*-test) and reliability of in-place field activity per lap (*p* = 0.014, Welch’s *t*-test) were significantly higher for dHPC^→NAc^ place cells compared to dHPC^−^ neurons (Figure 2G-H). These results suggest that the dHPC routes enhanced and more reliable spatial information to NAc compared to the general dHPC population, in line with previous results pointing towards the necessity of dHPC^→NAc^ projections for spatial memory expression (31, 32).

**Figure 2.**
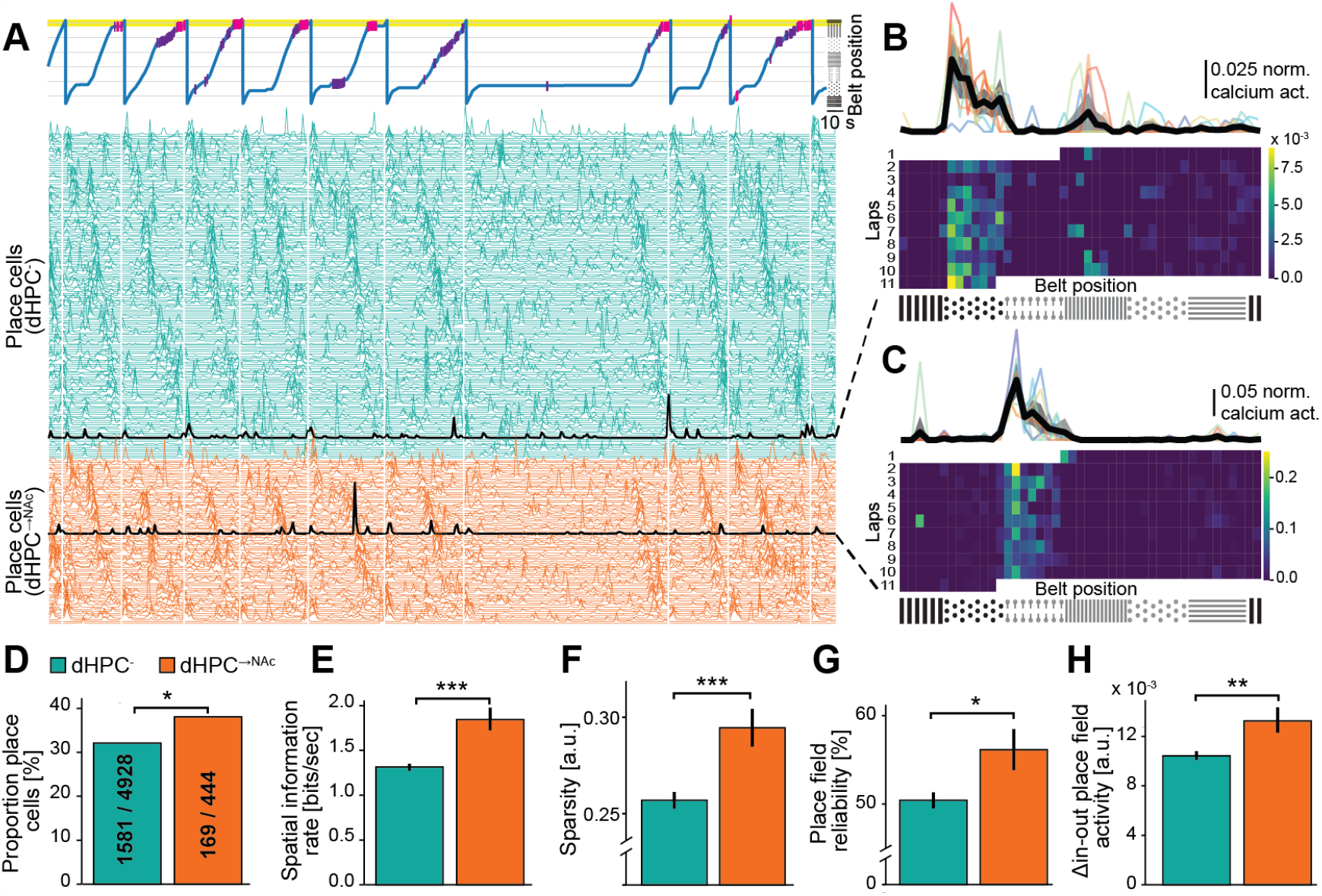
dHPC^→NAc^ neurons carry enhanced and more reliable spatial information. (A) Excerpt from one representative recording session showing behavioral activity (top) and denoised neural activity of identified place cells (bottom) over the course of several minutes from one mouse. White vertical lines mark new laps. Traces are ordered according to whether neurons co-expressed mCherry (red; dHPC^→NAc^) or not (green; dHPC^−^) and their respective place fields. Black traces are example neurons shown in (B and C). (B and C) Spatial activity of one neuron without (B) and with (C) mCherry co-expression over one session. Calcium events are binned by position on the belt (x axis) and laps (y axis) and show consistent activity at one position on the belt. (D-H) Comparisons of spatial tuning characteristics between dHPC^−^ (green) and dHPC^→NAc^ (red) neurons. NAc-projecting neurons contain a higher proportion of place cells (D, χ^2^(1, 5372) = 6.364); these place cells contain more spatial information per second (E, Welch’s *t* (186.55) = 4.770), show increased sparsity (F, Welch’s *t* (218.79) = 3.657), higher reliability (G, probability of firing maximally within their place field per lap; Welch’s *t* (203.46) = 2.479), and higher in-place field activity (H, Welch’s *t* (190.32) = 2.884). n = 6 mice, 19 imaging sessions, 5,372 (inc. 444 mCherry-coexpressing) neurons, 1,750 (inc. 169 mCherry-coexpressing) place cells. \**p* < 0.05, \*\**p* < 0.01, \*\*\**p* < 0.001. See also Figure S3.

### Place fields are modulated by local cue boundaries and are overrepresented near the reward zone

Previous studies found that place fields are often not homogenously distributed across the environment but can be modulated by salient environmental features such as textures, borders, or reward/goal zones (11, 49, 50). We hypothesized that information about such spatial features may be preferentially routed to NAc, given the relevance of NAc for spatial memory (29, 30) and particularly for reward-related behaviors (51).

Upon inspection of place cells’ spatially binned calcium profiles (Figure 3A-B and Figures S3, S4A), we noticed an over-abundance of place fields that seemed to cover entire texture zones, starting at the beginning of a texture zone and ending before the next texture zone began. To quantify this observation, we plotted the densities of place field start and end positions across the belt and compared the observed densities with those of randomly shuffled place fields. We found a significant overrepresentation of place field start and end positions (but not centers) near texture boundaries for both dHPC^−^ and dHPC^→NAc^ populations (ratios >99.9th percentile, permutation test). This effect was more pronounced in dHPC^→NAc^ neurons (PF start: *p* = 0.033, PF end: *p* = 0.036, χ^2^; Figure 3C-D and Figure S4A-C). This suggests that dHPC^→NAc^ neurons are more strongly modulated by local cue boundaries.

**Figure 3.**
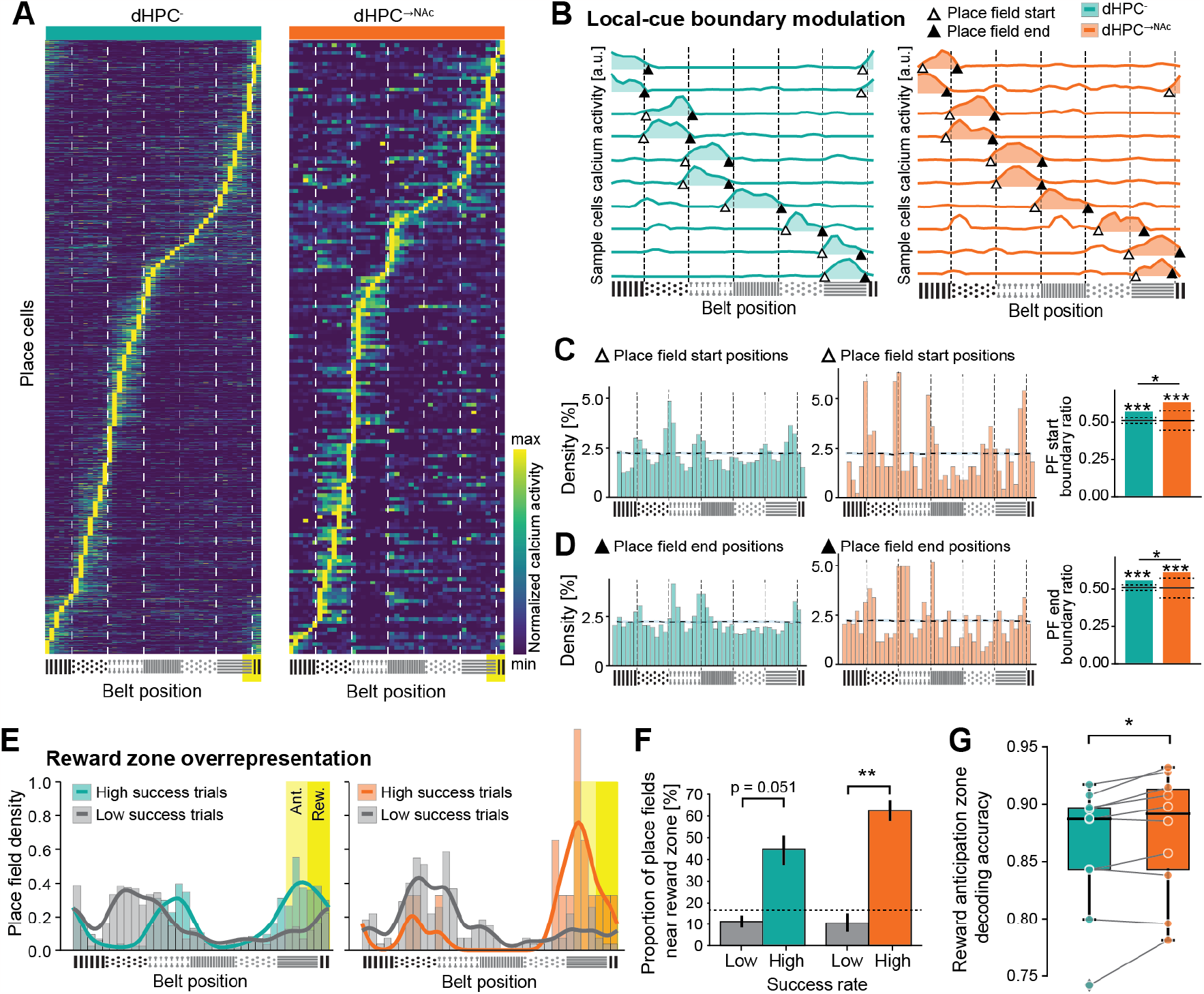
dHPC^→NAc^ place fields are modulated by local cues and reward zone. (A) Heat maps of dHPC^−^ (left) and dHPC^→NAc^ (right) place cells’ normalized position-binned average calcium events, ordered by place field location. Texture boundaries are indicated as white dashed lines. Yellow rectangle represents reward zone. (B Example place fields with edges near belt texture boundaries. Primary place fields are indicated by color fill. Triangles mark start (no fill) and end (fill) points. Dashed black lines represent texture boundaries. (C and D) Place field edges accumulate near texture boundary areas. Histograms of dHPC^−^ (green) and dHPC^→NAc^ (red) neurons’ place field start (C) and end (D). Dotted line and shade represent average and 95th CI of 1000x randomly shuffled place fields. Both dHPC^−^ and dHPC^→NAc^ place field start and end positions are significantly overrepresented at the 99.9th percentile (dotted black lines) compared to a randomly shuffled distribution. Start (χ^2^(1, 5134) = 5.735) and end positions (χ^2^(1, 5217) = 4.397) of dHPC^→NAc^ place fields are furthermore significantly overrepresented compared to the dHPC^−^ population. (E and F) Place cells are overrepresented near reward zone in high success trials. (E) Histograms (bars) and kernel density estimations (KDEs; lines) of place field centers for dHPC^−^ (left) and dHPC^→NAc^ (right) neurons, split into high success trials (green/red) and low success trials (gray). Reward zone (Rew.; yellow) and anticipation zone (Ant.; bright yellow) are indicated as rectangles. (F) Proportion of place fields in reward and vicinity zone is significantly higher in high-success trials (colored bars) compared to low-success trials (gray bars) in NAc-projecting neurons (red) but not in dHPC^−^ neurons (green). 2-way ANOVA, *F* _success_(1,1) = 54.918, *p* < 0.001, *F* _projection_(1,1) = 0.958, *p* = 0.338, *F* _interaction_(1,1) = 2.969, *p* = 0.098. Post-hoc Welch’s *t* -tests with Bonferroni correction: *t* _dHPC–_ (3.561) = 3.698; *t* _dHPC→NAc_ (3.479) = 8.671; *t* _low success_(13.629) = 0.093, *p* = 1; *t* _high success_(3.952) = 2.075, *p* = 0.215. Dashed line represents an even distribution of reward and anticipation zone. (G) A linear classifier shows significantly increased decoding accuracy of reward anticipation zone based on dHPC^→NAc^ neural activity compared to that of sample size-matched dHPC^−^ neurons. Wilcoxon’s *t* -test, *W* (9) = 5.0, *n* = 10 imaging sessions. All data are presented as mean ± SEM. \**p* < 0.05, \*\**p* < 0.01, \*\*\**p* < 0.001. See also Figure S4.

Mice were trained to lick for reward in a hidden reward zone. Previous studies have shown that such zones and their preceding vicinity are often over-represented by place cells (35, 52, 53) – we hypothesized that this effect might involve dHPC^→NAc^ neurons, given the NAc’s role in reward-related behaviors (51). When pooled across all sessions, we found little evidence of such an overrepresentation (Figure S4D-E). Mouse behavior generally shows great variability though, and previous work demonstrated the dependence of such an overrepresentation on individual task success (35, 52). We thus divided sessions into high- and low-success based on lick performance (lick precision and reward dispensation, see Methods) and found significantly more dHPC^→NAc^ place cells near the reward zone (reward and anticipation zones) in high-success sessions compared to low-success sessions (*p* = 0.0036, Welch’s *t*-test). In contrast, dHPC^−^ neurons only showed a trend towards significance (*p* = 0.051, Welch’s *t*-test; Figure 3E-F). In line with previous studies, both neuronal populations also showed a strong correlation between the success rate (percentage of rewarded laps) and the proportion of place fields near the reward zone across sessions (*r* = 0.62; Figure S4F).

If reward zone information is preferentially encoded in dHPC^→NAc^ populations and NAc neurons play an integral part in reward-related behaviors, we would expect linear decoders to identify reward or anticipation zones more accurately based on dHPC^→NAc^ activity compared to that of dHPC^−^. To test this, we trained a linear classifier based on a support vector machine (SVM) with calcium activity and reward zone information on odd/even laps and tested decoding accuracy on even/odd laps, using either dHPC^−^ or sample size-matched dHPC^→NAc^ populations (see Methods). We found that, within individual sessions, decoding accuracy of the reward anticipation zone was significantly enhanced for dHPC^→NAc^ populations (*p* = 0.0195, Wilcoxon’s test; Figure 3G). These findings demonstrate significant modulation of dHPC^→NAc^ neurons by local cue boundaries and enhanced reward zone coding.

### Enhanced coding of low velocities in dHPC^→NAc^ neurons

As correct performance in the spatial reward learning task goes hand in hand with a reduction in velocity and an increase of licking near the reward zone (see Figure S1K-N), we wondered if such non-spatial task-relevant behavioral features were encoded by dHPC neurons and its projections to NAc (9, 54, 55). We hypothesized that NAc may have privileged access to information on low velocities as mice generally slow down near the reward zone, presumably to allow for better discrimination and to engage in anticipatory licking. In line with previous analyses of speed coding in hippocampal and parahippocampal regions (8, 56), we av-eraged each neuron’s calcium activity per velocity bin from 2 to 30 cm/s and regressed this activity against velocity. Neurons with a significant regression model (after correcting for false discovery rate) and positive slope were classified as speed-excited (Figure 4A-B). We found around 15 % of neurons were positively speed-modulated, with comparable proportions between dHPC^−^ and dHPC^→NAc^ neurons (13 % vs. 16 %, *p* = 0.109, χ^2^; Figure 4C-D). We also identified neurons with a significant velocity regression but a negative slope (Figure 4E-F). We found approximately 15 % of such speed-inhibited neurons, with a significantly larger proportion among the dHPC^→NAc^ population (15 % vs. 21 %, *p* = 0.00022, χ^2^; Figure 4G-H). These results suggest widespread modulation of dHPC neuronal activity by non-spatial features such as velocity, with dHPC^→NAc^ neurons specifically over-representing low speeds.

**Figure 4.**
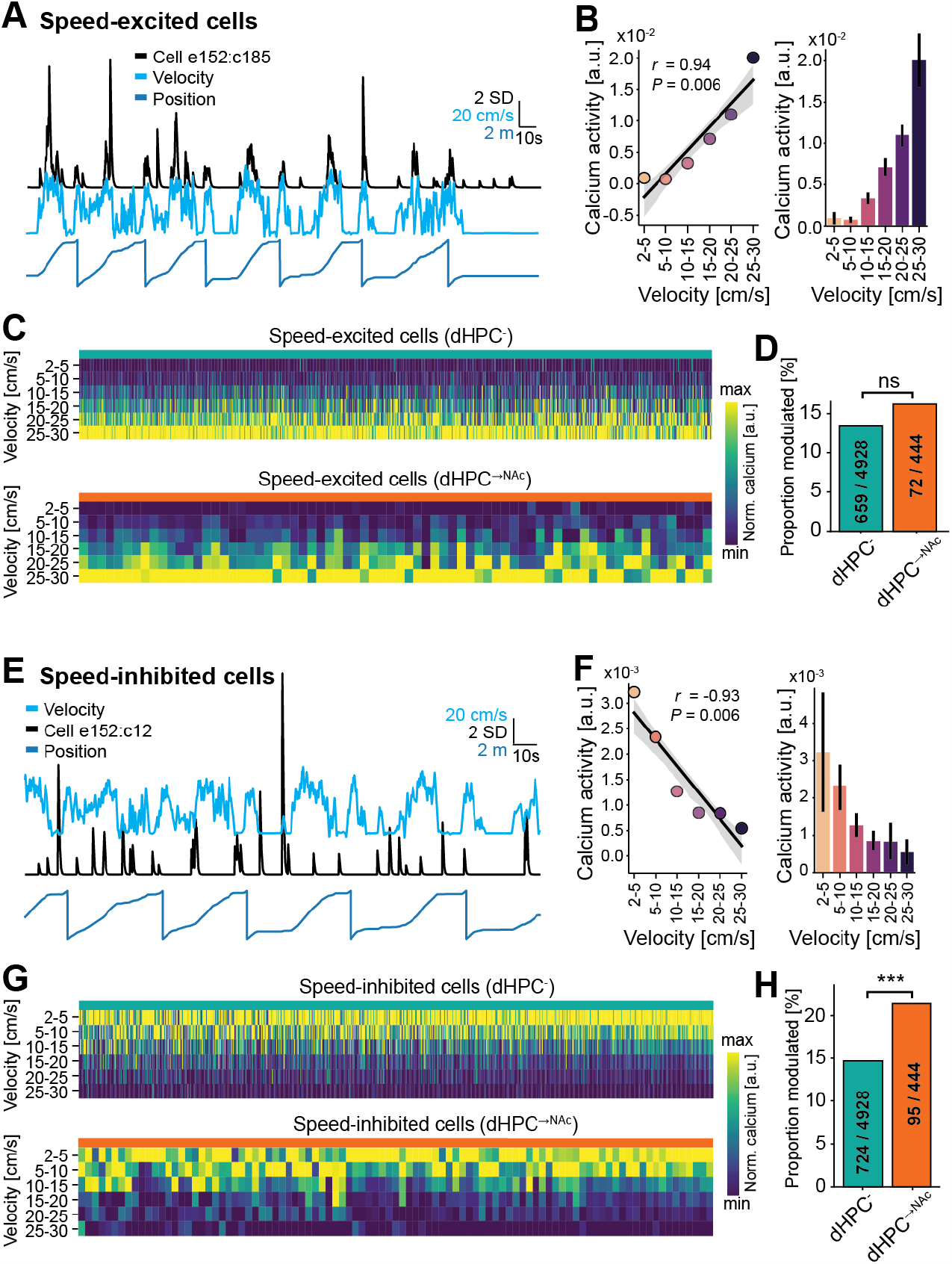
Speed-inhibited cells are overrepresented in dHPC^→NAc^ neurons. (A-D) Speed-excited dHPC neurons. (A and B) Representative example of one speed-excited neuron. (A) Sample traces of velocity, position and one neuron’s denoised calcium activity. Note the increased calcium activity in times of high velocity, irrespective of position. (B) Linear regression on average calcium events per velocity bin shows a significant positive relationship (slope = 7.43*×*10^−4^, intercept = -2.097*×*10^−3^, *r* = 0.937, *p* = 0.0058). (C) Heatmaps of speed-binned normalized calcium activity of all significantly positively speed-modulated dHPC^−^ (top) and dHPC^→NAc^ (bottom) neurons. (D) Proportions of speed-excited neurons are comparable between dHPC^−^ and dHPC^→NAc^ populations (χ^2^(1, 5372) = 2.565, *p* = 0.109). (E-H) Speed-inhibited dHPC neurons. (E and F) Representative example of one speed-inhibited neuron. (E) Sample traces of velocity, position and one neuron’s denoised calcium activity. Note the increased calcium activity in times of low velocity, irrespective of position. (F) Linear regression on average calcium events per velocity bin shows a significant negative relationship (slope = -1.0437*×*10^−4^, intercept = 2.814*×*10^−3^, *r* = -0.933, *p* = 0.0065). (G) Heatmaps of speed-binned normalized calcium activity of all significantly negatively speed-modulated dHPC^−^ (top) and dHPC^→NAc^ (bottom) neurons. (H) Negatively tuned neurons are overrepresented in the NAc-projecting population (χ^2^(1, 5372) = 13.66, *p* = 0.00022). Speed-modulated neurons were classified as showing a significant linear regression at *p* < 0.05 after Benjamini/Hochberg FDR correction. All data are presented as mean ± SEM. ns: not significant, \*\*\**p* < 0.001.

### Overrepresentation of appetitive licking-excited dHPC^→NAc^ neurons

Besides a decrease in velocity when approaching the reward zone, mice also increasingly engaged in licking behavior (see Figure 1B-D). Given the NAc’s dual role in appetitive and consummatory behaviors (57), we wondered if licking behaviors might be reflected in the neural activity of dHPC^→NAc^ neurons. For this, we distinguished between consummatory licking which occurs after a reward is dispensed and allows the mouse to consume the reward provided, and appetitive licking which is an operant behavior that will lead to reward dispensation when performed at the correct location on the belt. We found a significant decrease of calcium activity during reward consumption in both dHPC^−^ and dHPC^→NAc^ populations (Figure S5A-C).

Appetitive licking, on the other hand, had no apparent effect on neural activity in dHPC^−^ neurons but coincided with a significant increase in calcium activity in the dHPC^→NAc^ population that began about one second before lick onset and then fell below baseline 4 seconds after licking (Figure 5A-C). We investigated if this population-averaged data is reflected on the single-cell level and if there are individual cells that are reliably modulated by appetitive licking (Figure 5D-E and Figure S5D-E). Comparing the pre- and post-lick neural activity for each neuron for each appetitive lick event, we identified a total of 1,268 neurons (24 %) that were significantly (negatively or positively) modulated by appetitive licking (Figure 5F-I). Interestingly, while the proportion of lick-inhibited neurons was comparable between dHPC^−^ and dHPC^→NAc^ populations (19.7 % vs 17.1 %, *p* = 0.20, χ^2^; Figure 5I), we found a significantly larger proportion of lick-excited neurons in dHPC^→NAc^ populations (3.8 % vs 7.4 %, *p* < 0.001, χ^2^; Figure 5H). These findings suggest that dHPC does not route reward information per se to NAc, but rather information on appetitive behaviors required to obtain such rewards.

**Figure 5.**
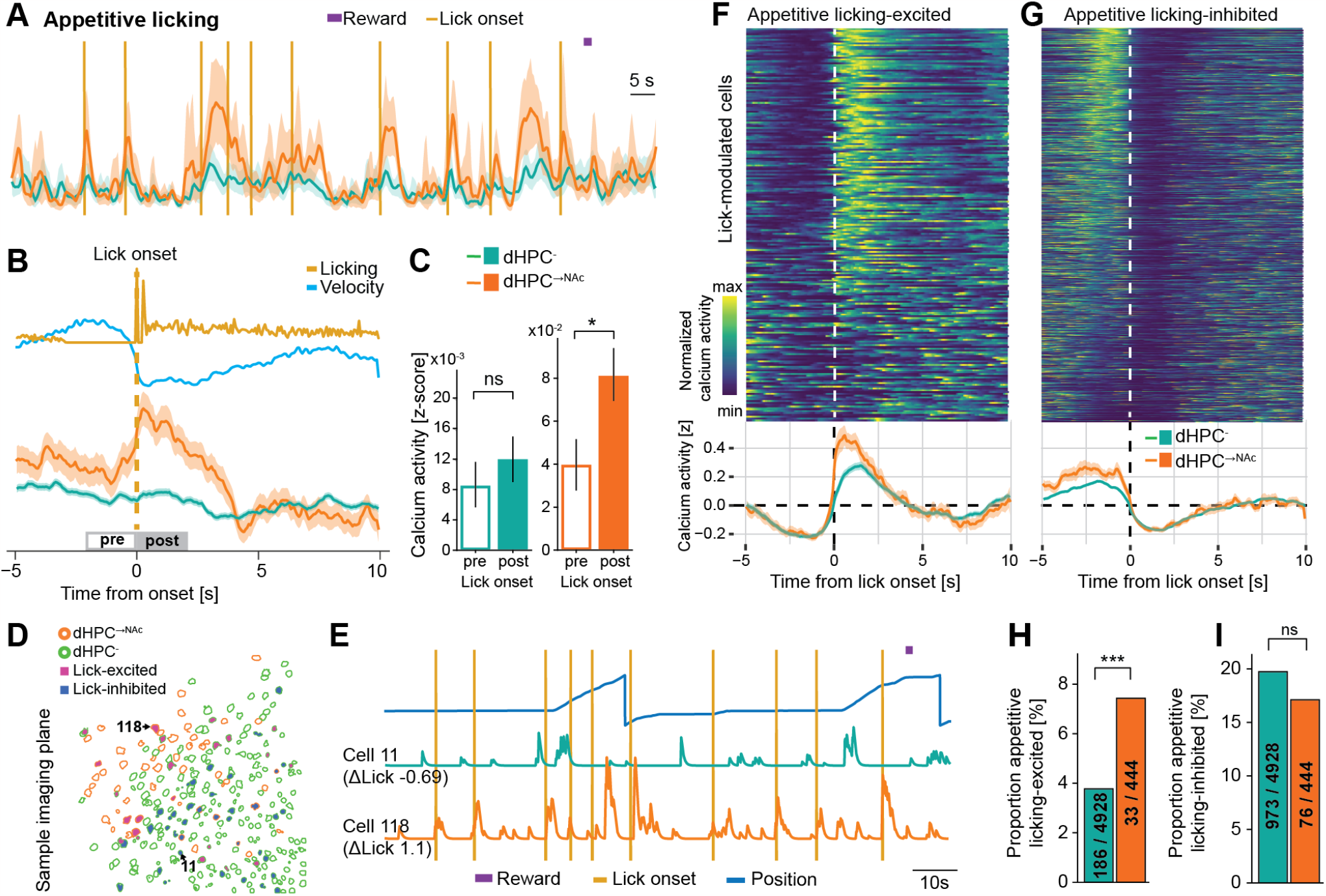
dHPC^→NAc^ neurons are excited by appetitive licking and are over-represented in lick-excited neurons. (A-C) Appetitive licking is accompanied by increased neural activity in dHPC^→NAc^ neurons but not dHPC^−^ neurons. (A) Representative example showing reward dispensation (purple bar) and appetitive licking onsets (golden vertical lines) and population calcium activity from dHPC^−^ (green) and dHPC^→NAc^ (red) neurons. Note the robust increase of neural activity around lick onsets in the dHPC^→NAc^ population. (B) Event-triggered average traces around appetitive licking onset, including licking and speed as well as population average calcium activity of dHPC^−^ (green) and dHPC^→NAc^ (red) neurons. Gray rectangles indicate time windows for comparing calcium activity shown in (C). (C) Calcium activity is differentially modulated by appetitive licking onset only in dHPC^→NAc^ neurons; two-way mixed ANOVA; *F* _licktiming_(1, 5370) = 2.843, *p* = 0.0918, *F* _projection_(1, 5370) = 43.779, *p* < 0.001 *F* _interaction_(1, 5370) = 7.073, *p* = 0.0079. Post-hoc *t* -tests with Bonferroni correction: t_dHPC–_ (4927) = 0.871, *p* = 0.768; t_dHPC→NAc_ (443) = 2.470, *p* = 0.0277. (D and E) Example imaging session and traces showing lick-excited and lick-inhibited neurons. (D) Field of view showing spatial profiles of dHPC^−^ (green outlines) and dHPC^→NAc^ (red outlines), some of which are classified as lick-excited (violet fill) or lick-inhibited (dark blue fill); neurons #11 and #118 are highlighted. (E) Behavioral traces and calcium activity of sample neurons #11 (lick-inhibited) and #118 (lick-excited). (F and G) Heatmaps (top) and event-triggered calcium activity averages (bottom) of neurons classified as appetitive licking-excited (F) and licking-inhibited (G). (H) Proportion of appetitive licking-excited neurons is higher in NAc-projecting neurons; χ^2^(1, 5372) = 13.018, *p*= 0.00031. (I) Proportion of appetitive licking-inhibited neurons is not different between populations; χ^2^(1, 5372) = 1.626, *p* = 0.202. All data are presented as mean ± SEM. ns: not significant, \**p* < 0.05, \*\*\**p* < 0.001. See also Figure S5.

### Optogenetic activation of dHPC terminals in NAc induces mouth movement and deceleration

Given the NAc’s hypothesized role as an interface between limbic and motor systems (25), we investigated if the observed increase of calcium activity in dHPC^→NAc^ neurons may be a corollary signal of motor action or indeed serve a causal role in appetitive behaviors (58). To enable high-resolution behavioral tracking, we monitored mouse orofacial movements using a high-speed near-infrared camera (Figure 6A-C). To test for a causal role of excitatory dHPC^→NAc^ projections in appetitive behaviors, we injected animals with either CaMKIIa-driven ChR2 or EYFP into dHPC and implanted light fibers in the NAc (*n* = 4/3 mice; Figure 6D-F). After habituating mice to run on the treadmill and receive rewards upon licking on the lick spout, mice were given 5 mW of 473 nm 20 Hz (5 ms duration) pulsed laser light for up to 10 seconds upon entry into a hidden light stimulation zone.

**Figure 6.**
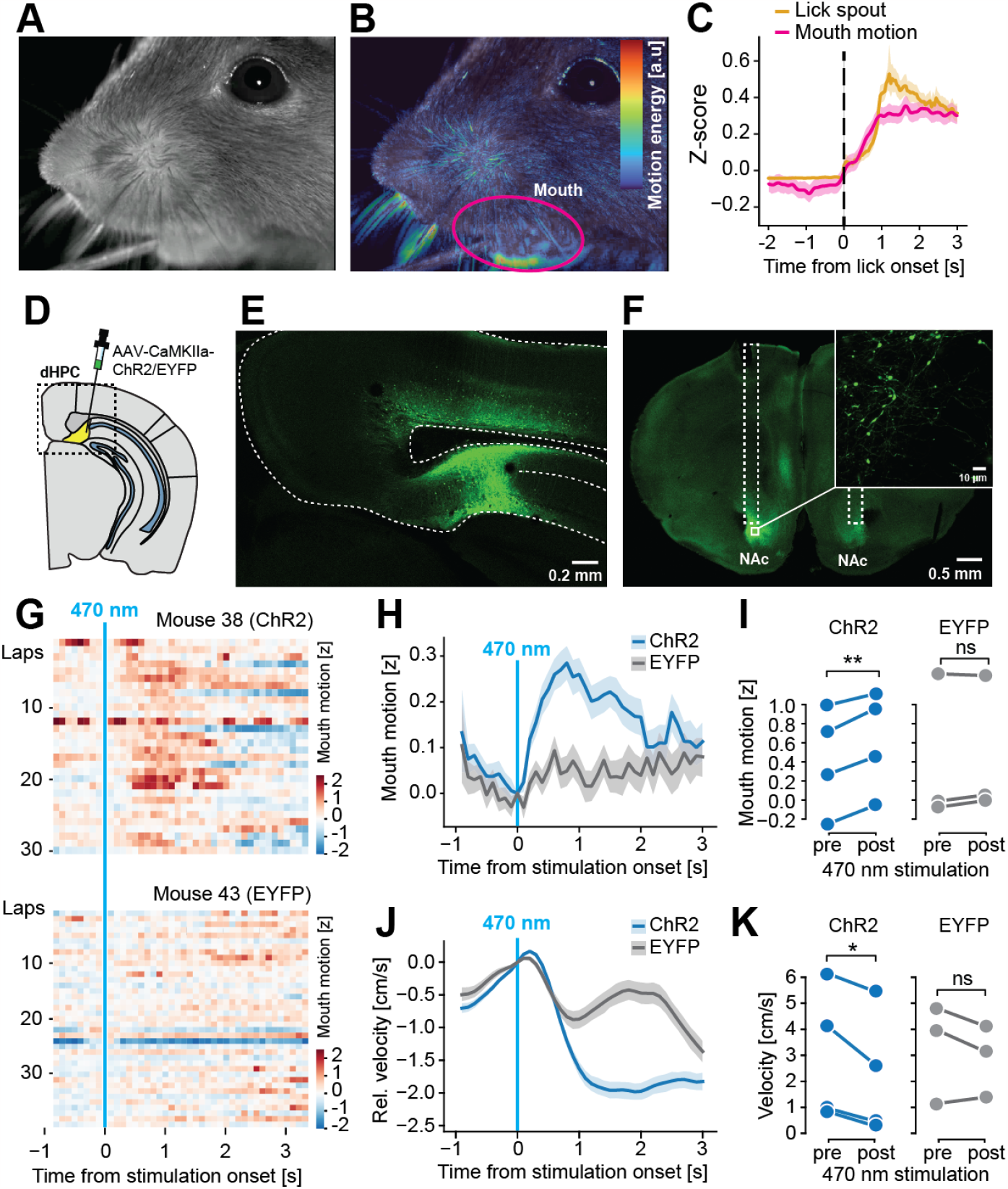
Optogenetic stimulation of dHPC axons in NAc induces mouth movement and deceleration. (A-C) Tracking of orofacial movements via infrared camera recordings. (A) Example still image from near-infrared camera, sampled at 75 Hz. (B) False-color coded motion energy (pixel-by-pixel intensity difference to previous image) overlaid over sample image from (A). Automatically segmented mouth region is indicated by a purple line. (C) Average motion energy in mouth region increases with licking. Golden line shows analog lick spout signal. (D-F) Experimental approach for optogenetic dHPC^→NAc^ stimulation. (D) Injection schematic. (E) Somatic expression of ChR2-EYFP in dorsal pro-subiculum. (F) Axonal expression of ChR2-EYFP in the NAc, where light fibers are placed (tracts indicated by white dotted lines). (G) Representative examples of mouth motion around onset of optogenetic stimulation in an animal expressing ChR2 (top) or EYFP control (bottom). (H) Trial-averaged mouth motion activity around time of optogenetic stimulation in ChR2 (blue) and EYFP (gray)-expressing mice. Mouth motion is significantly increased with optogenetic stimulation in ChR2 animals (*t* (3) = 7.485; *p* = 0.00494) but not EFYP animals (*t* (2) = 1.353; *p* = 0.309). Paired *t* -tests; *n* = 4 mice (ChR2), *n* = 3 mice (EYFP). (J) Trial-averaged relative velocity around time of optogenetic stimulation in ChR2 (blue) and EYFP (gray)-expressing mice. (K) Velocity is significantly decreased with optogenetic stimulation in ChR2 animals (*t* (3) = -3.551; *p* = 0.0381) but not EFYP animals *t* (2) = -1.263; *p* = 0.334. Paired *t* - tests; *n* = 4 mice (ChR2), *n* = 3 mice (EYFP). All data are presented as mean ± SEM. ns: not significant, \**p* < 0.05, \*\**p* < 0.01.

We found that, shortly after stimulation onset, ChR2-expressing mice reliably showed increased mouth movement for up to two seconds after stimulation, while we observed no effects in mice expressing EYFP (*p*_ChR2_ = 0.0099, *p*_EYFP_ = 0.617, paired *t*-tests; Figure 6G-H). In line with this, we also found a significant deceleration of running on the treadmill upon light delivery in ChR2 animals but not EYFP animals (*p*_ChR2_ = 0.0381, *p*_EYFP_ = 0.334, paired *t*-tests; Figure 6J-K). These findings support the idea that dHPC^→NAc^ projections may not simply represent task-related non-spatial behavioral features but may in fact represent a driving force in the generation of task-relevant appetitive behavior.

### Enhanced conjunctive coding of space, velocity, and appetitive behaviors in dHPC^→NAc^ neurons

We identified cells modulated by space, velocity, and appetitive licking. Previous studies suggested that individual hippocampal neurons do not necessarily exclusively code for one single feature but are instead able to conjunctively encode various environmental properties (9, 55, 59, 60). Such conjunctive coding may be particularly relevant for downstream linear decoders to select task-appropriate actions (61, 62). We thus investigated speed and lick modulation of projection-specific place cells and interactions between velocity and lick modulation.

We first analyzed speed coding in dHPC^→NAc^ place cells and compared it to the dHPC^−^ population (Figure 7A-B). We found about one third of hippocampal place cells were also speed-inhibited, in contrast to only about 7 % of non-place cells. This effect was particularly pronounced in dHPC^→NAc^ place cells (43 % vs. 31 %, *p* = 0.0027, χ^2^). Conversely, place cells were significantly less likely to be speed-excited than non-place cells (10 % vs. 15 %, *p* < 0.001, χ^2^), an effect that was again more pronounced in dHPC^→NAc^ neurons. This shows that dHPC place cells, and in particular those projecting to NAc, are more likely to be speed-inhibited, and less likely to be speed-excited.

**Figure 7.**
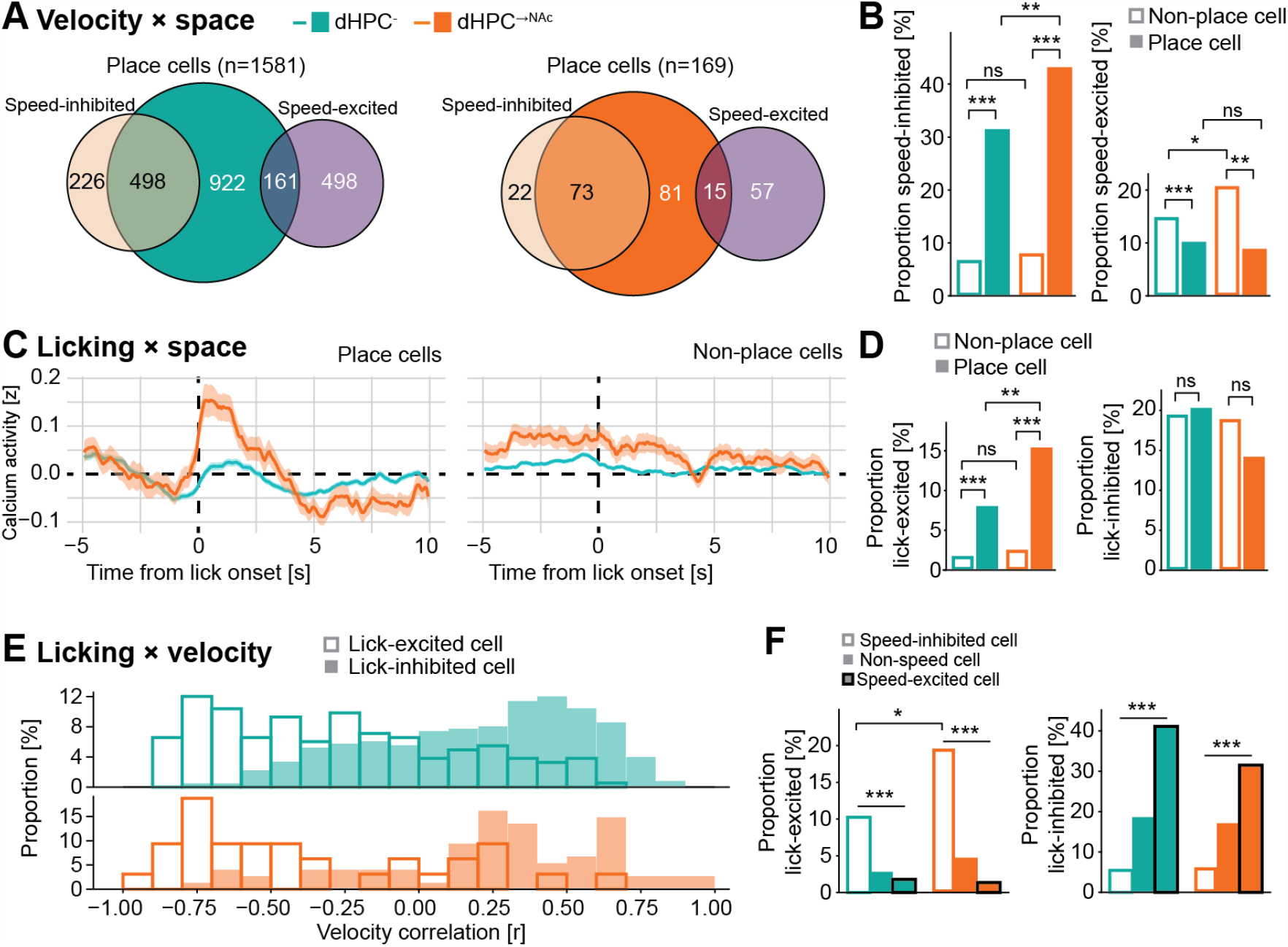
Enhanced conjunctive coding of space, velocity and licking in NAc-projecting neurons. (A and B) Conjunctive coding of space and velocity. (A) Venn diagrams of dHPC^−^ (left) and dHPC^→NAc^ (right) place cells and their overlaps with negatively (ocher) and positively (purple) tuned speed cells. Numbers shown refer to absolute numbers of neurons classified. (B) Proportions of dHPC^−^ (green) and dHPC^→NAc^ (red) place cells (fill) and non-place cells (no fill) that are also significantly speed-modulated. A larger proportion of place cells is speed-inhibited compared to non-place cells (dHPC^−^: χ^2^(1, 4928) = 522.71; dHPC^→NAc^: χ^2^(1, 444) = 75.02). dHPC^→NAc^ place cells also have a higher proportion of speed-inhibited cells than dHPC^−^ place cells (χ^2^(1, 1750) = 8.977). Speed-excited neurons are overrepresented in non-place cells compared to place cells (dHPC^−^: χ^2^(1, 4928) = 20.034; dHPC^→NAc^: χ^2^(1, 444) = 9.966). dHPC^→NAc^ non-place cells also have a higher proportion of speed-excited cells than dHPC^−^ non-place cells (χ^2^(1, 3622) = 6.255). (C and D) Conjunctive coding of space and licking. (C) Event-triggered average calcium traces for dHPC^−^ (green) and dHPC^→NAc^ (red) place cells (left) and non-place cells (right). (D) Proportions of lick-excited neurons are significantly enriched in dHPC^−^ (χ^2^(1, 4928) = 114.515) and dHPC^→NAc^ (χ^2^(1, 444) = 23.248) place cells compared to non-place cells. The proportion of dHPC^→NAc^ lick-excited place cells is also higher than the proportion of dHPC^−^ lick-excited place cells (χ^2^(1, 1750) = 9.442); there is no difference between non-place cells (χ^2^(1, 3622) = 0.488, *p* = 0.485). There is no difference in the proportions of lick-inhibited place and non-place dHPC^−^ and dHPC^→NAc^ neurons (χ^2^s, all *p* > 0.05). (E and F) Conjunctive coding of velocity and licking. (E) Histogram of lick-excited (no fill) and lick-inhibited (fill) cells’ velocity correlations (green: dHPC^−^; red: dHPC^→NAc^). Note the negative velocity correlations of lick-excited cells and positive velocity correlations for lick-inhibited cells. (F) Proportions of lick-excited (left) and lick-inhibited (right) cells among speed-inhibited (no fill), non-speed-modulated (fill) and speed-excited (fill, black stroke) dHPC^−^ (green) and dHPC^→NAc^ (red) cells. Proportions of lick-excited cells are overrepresented in speed-inhibited cells (dHPC^−^: χ^2^(2, 4928) = 100.484; dHPC^→NAc^: χ^2^(2, 444) = 27.608). Lick-excited neurons are further enriched in speed-inhibited dHPC^→NAc^ neurons compared to dHPC^−^ neurons (χ^2^(1, 830) = 6.564). Proportions of lick-inhibited cells are overrepresented in speed-excited cells (dHPC^−^: χ^2^(2, 4928) = 290.832; dHPC^→NAc^: χ^2^(2, 444) = 18.825). All data are presented as mean ± SEM. ns: not significant, \**p* < 0.05, \*\**p* < 0.01, \*\*\**p* < 0.001.

We next analyzed lick modulation of place cells and, surprisingly, found that the previously observed lick-related increase in calcium activity (Figure 5B) was largely carried by place cells and not by non-place cells (Figure 7C). This effect seems to be mostly carried by lick-excited neurons that are significantly overrepresented in place cells compared to non-place cells (8 % vs. 2 %, *p* < 0.001, χ^2^), particularly in dHPC^→NAc^ neurons (15 % vs. 3 %, *p* < 0.001, χ^2^; Figure 7D). Lick-inhibited neurons, on the other hand, were distributed equally between place and non-place dHPC^−^ and dHPC^→NAc^ neurons. These findings show that a large majority of lick-excited neurons, especially in the dHPC^→NAc^ population, also code for spatial information. Finally, we investigated interactions between lick and velocity modulation. Comparing velocity correlations of lick-excited and lick-inhibited neurons, we observed a clear skew of lick-excited cells to have more negative velocity correlations and lick-inhibited cells to have more positive velocity correlations, visible in both dHPC^−^ and dHPC^→NAc^ populations (Figure 7E). This results in significantly more speed-inhibited cells to be lick-excited compared to speed-excited cells (11 % vs. 2 %, *p* < 0.001, χ^2^), an effect that was even more pronounced in dHPC^→NAc^ neurons (20 % vs. 1 %, *p* < 0.001, χ^2^; Figure 7F). Conversely, speed-excited neurons were much more likely to be lick-inhibited than speed-inhibited neurons (40 % vs. 6 %, *p* < 0.001, χ^2^). These results demonstrate that there is a strong inverse relationship between lick and velocity modulation.

One caveat of such conjunctive coding analyses is that in our behavioral task, trained mice often show highly stereotypical behavior, such that mice would mostly lick at one location where they would also slow down (see Figure 1C and Figure S1K-N). In light of this, conjunctive coding could be an epiphenomenon of collinear behavioral features. To account for this collinearity, we modelled the influence of three key behavioral features (space, velocity, and appetitive licking) on the activity of each neuron by building a generalized linear model for each neuron (GLM; Figure 8A). On average, we found that our models could explain close to 40% of the variance observed in our test datasets (Figure S6A), with dHPC^→NAc^ neurons showing increased feature importance for position and licking (Figure S6B). To determine significant contributions of the three behavioral features, we next built 3 × 100 models in which one of the behavioral features was randomly shuffled against time, and compared the variance explained to the original model (Figure 8A; see also ref (63)). Thus, if neural activity spuriously coincided with the activity of one behavioral feature that could be similarly explained by a largely collinear behavioral feature, the model’s predictive performance should be mostly unaffected by this shuffling. The average drop in variance explained to the full model was significantly stronger in dHPC^→NAc^ neurons compared to dHPC^−^ for position and velocity but not licking (Figure S6C). We classified significant modulation as behavioral features whose shuffling led to a reduction in variance explained in more than 95% of shuffled models. We found that cells thus encoding space, velocity, and licking were overrepresented in dHPC^→NAc^ compared to dHPC^−^ neurons (space: *p* < 0.001; velocity: *p* < 0.001; licking: *p* = 0.0015; χ^2^; Figure 8B). Importantly, we also found significantly increased proportions of conjunctive coding for all three feature combinations as well as triple-conjunctive neurons in dHPC^→NAc^ neurons compared to dHPC^−^ (all combinations *p* < 0.001, χ^2^; Figure 8C-D). This results in a significantly higher proportion of conjunctive coding neurons in dHPC^→NAc^ neurons compared to dHPC^−^ (44 % vs. 19 %, *p* < 0.001, χ^2^; Figure 8E).

**Figure 8.**
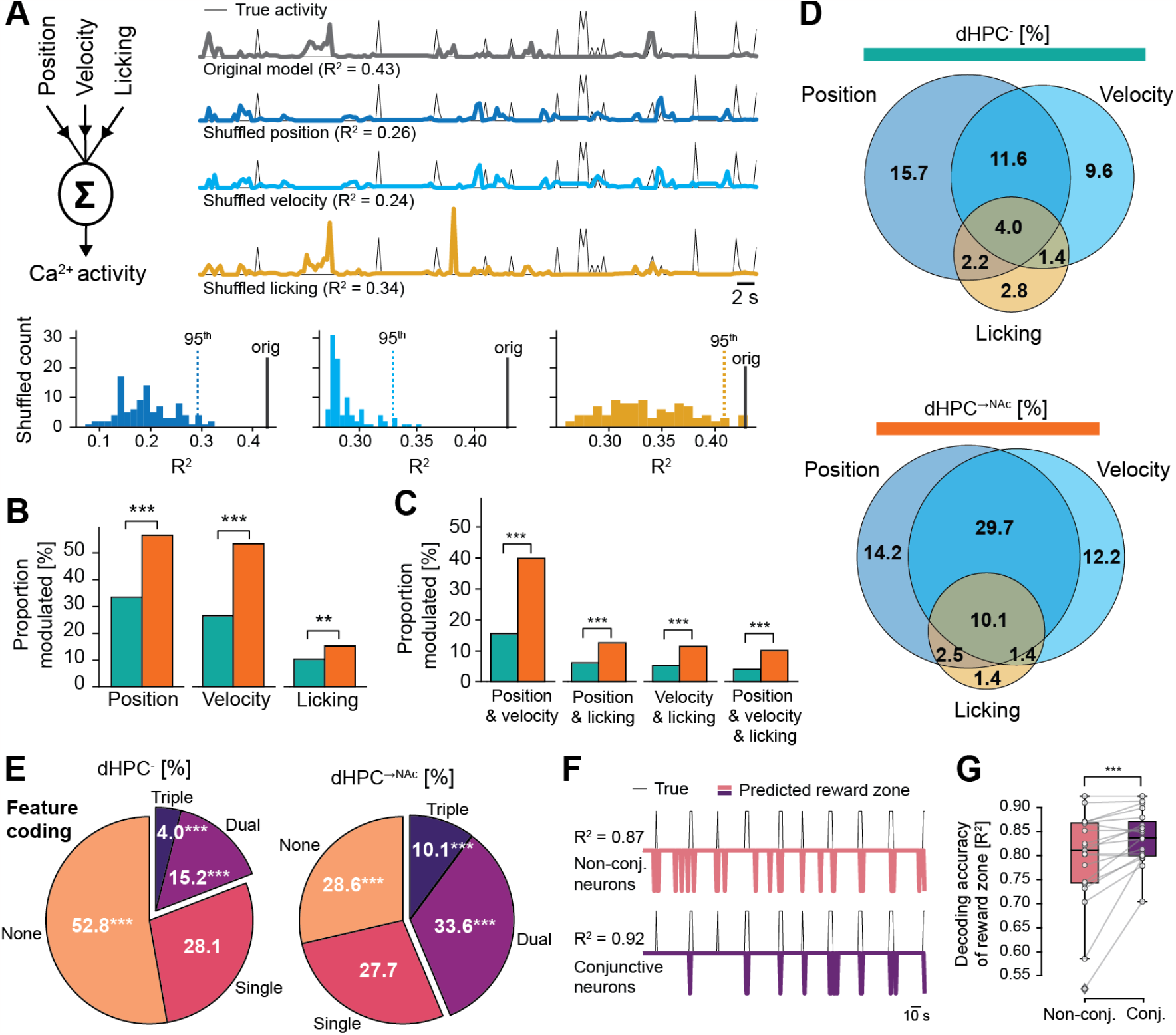
A generalized linear model confirms enhanced conjunctive coding in NAc-projecting neurons. (A) Schematic of the generalized linear model (GLM) using position, velocity, and appetitive licking data as predictors for each neuron’s calcium activity (left). Example modelling approach for a triple conjunctive neuron (right). Upper traces show true calcium activity (downsampled and normalized) of test dataset (thin black line) as well as predictions of original model (thick gray) and one example each for each shuffled feature. Note the reductions in prediction quality when features are shuffled. Bottom histograms show *R*^2^ distributions for 100 shuffled models for each feature. Dotted line represents value of 95th percentile, thick black line represents original model’s *R*^2^. (B) Increased proportions of dHPC^→NAc^ neurons modulated by position (χ^2^(1, 5372) = 93.634), velocity (χ^2^(1, 5372) = 141.86), and licking (χ^2^(1, 5372) = 10.050). (C) Increased proportions of conjunctive coding in dHPC^→NAc^ neurons for position & velocity (χ^2^(1, 5372) = 163.97), position & licking (χ^2^(1, 5372) = 26.029), velocity & licking (χ^2^(1, 5372) = 27.145), and position & velocity & licking (χ^2^(1, 5372) = 34.993). (D) Venn diagrams showing overlap of neurons GLM-classified as modulated by position, velocity and licking in dHPC^−^ (top) and dHPC^→NAc^ (bottom) neurons. Proportions of *n*-feature coding neurons in dHPC^−^ (left) and dHPC^→NAc^ (right) populations. Non-coding neurons are overrepresented in dHPC^−^ neurons (χ^2^(1, 5372) = 35.382); single-coding neurons are comparably distributed (χ^2^(1, 5372) = 0.0057); dual-coding (χ^2^(1, 5372) = 61.336) and triple-coding (χ^2^(1, 5372) = 30.447) neurons are overrepresented in dHPC^→NAc^ neurons. (F and G) A linear classifier to decode presence of reward zone. (F) Example true presence of reward zone (thin line) and decoder predictions based on non-conjunctive neurons (pink; top) and conjunctive-coding neurons (dark purple; bottom). (G) Conjunctive-coding neurons allow a linear decoder to classify the presence of reward zone more accurately than non-conjunctive coding neurons (Wilcoxon’s *W* (18) = 14.0). \*\**p* < 0.01, \*\*\**p* < 0.001. See also Figure S6.

As conjunctive coding has been suggested to aid downstream linear decoders to select task-appropriate actions (55, 61), we wondered if this increased conjunctive code might allow linear decoders to identify the presence of the reward zone more correctly in our task. We thus trained an SVM-based linear classifier on each trial’s odd/even laps’ reward zones and tested the decoding accuracy on even/odd laps, based on conjunctive or randomly sample size-matched non-conjunctive coding neurons (Figure 8F). We found enhanced reward zone decoding accuracy for conjunctive compared to non-conjunctive coding neurons (*p* < 0.001, Wilcoxon’s test; Figure 8G). Thus, our data suggest that the enhanced conjunctive coding observed in dHPC^→NAc^ neurons allows NAc neurons to better identify the reward zone and guide task-appropriate appetitive behavior.

## Discussion

While our understanding of the principal mechanisms by which internal and environmental features are processed by hippocampal circuits has greatly advanced (5, 6), the information content of its various output streams and their specific contributions to behavior remain largely unresolved. The NAc, with its proposed role as an integrator between limbic and motor systems (25) seems an ideal candidate to transform hippocampal mnemonic and contextual information into task-relevant behaviors (26), but evidence for such a role has remained sparse.

One main finding was the identification of a larger proportion of place cells among the dHPC^→NAc^ population that also encoded more spatial information and showed both enhanced trial-to-trial reliability and in-place-field activity. Our data confirm previous suggestions that the NAc receives spatial information from dHPC (26) and likely explains previously described spatial tuning of NAc neurons (29). The dHPC^→NAc^ population’s observed enhanced spatial tuning and stronger modulation of local cues parallels previously described differences between deep and superficial dHPC neurons (18). Indeed, vHPC → NAc projection neurons have been shown to accumulate mostly in deep layers (64), raising the possibility that previously observed anatomical-functional differences could at least partly be explained by differences in projection targets, thus adding another important layer of complexity to pyramidal cell heterogeneity (17).

Place cells in CA1 and subiculum have long been appreciated to overrepresent reward/goal zones (53, 65, 66) in a way that correlates with behavioral performance (35, 52). This effect likely depends on the interaction of inputs from entorhinal cortex layer 3, dopaminergic inputs from VTA, and hippocampal NMDA receptors (36, 52, 67). While previous studies suggested a role for a largely undefined dedicated population of dHPC cells in this overrepresentation (65) and differences have been observed along the radial dCA1 axis (35), evidence for projection-specific reward zone representations has remained sparse. Although synchronous activity has been observed between dHPC and NAc during spatial reward learning tasks (38, 68, 69), in line with a central role of NAc in reward learning (reviewed in ref (70)), our findings present direct evidence of increased reward zone overrepresentation by dHPC^→NAc^ neurons. Interestingly, a previous study (19) showed projection-specific ambiguity of reward zone activity depending on vCA1 → NAc collaterals. Future studies should investigate potential functional differences in collaterals of dHPC projections (41).

Successful behavioral performance in our spatial reward learning task depended on a decrease in velocity and increased appetitive lick activity as mice approached the hidden reward zone. In line with previous work (8, 9, 12, 20), we found a considerable proportion of speed-modulated dHPC neurons. Interestingly, NAc-projecting neurons were more likely to be speed-inhibited, suggesting an elevated role in reward approach behaviors (see also ref (12)). Similarly, we observed significant modulation of hippocampal neurons during appetitive and consummatory licking (see also refs (12, 54)). This modulation was dichotomous: consummatory licking resulted in widespread suppression of dHPC activity, while appetitive licking led to enhanced activity in dHPC^→NAc^ neurons. Activity of dHPC neurons, and especially those projecting to NAc, thus seems to mirror the activity within NAc: Inhibition of NAc D1R medium spiny neurons (MSNs) has been observed during lick behavior (71–73) and was shown to be both necessary (71, 74) and sufficient (71, 75) for consummatory licking. We found that the origin of these signals may be localized upstream, similar to recent findings of an inhibitory permissive drive for licking in vHPC projections to the NAc (72, 76) and that inhibiting these projections facilitates licking (72). This suggests a similar involvement of dHPC^−^ and vHPC-accumbens projections for consummatory licking.

Contrasting this consumption-related inhibition, we found appetitive lick-related dHPC^→NAc^ excitation and optogenetic induction of lick-related behaviors. This dichotomy likely reflects previously described differences in the NAc’s role for consummatory and appetitive behaviors (57): vHPC → NAc activity was shown to increase during active food investigation (76) and around the time of lever pressing to obtain a reward (Iyer E, Muir J, Namuhoranye B, Bagot, R. Glutamatergic afferents to the nucleus accumbens integrate outcomes in reward-learning. Poster S01-181, FENS Forum 2022), while optogenetic stimulation of this projection facilitated nose poke and lever pressing behaviors (33, 77). Given that optogenetic stimulation of D2R MSNs in the NAc shell was also found to produce repetitive jaw movements (78), it is likely that D1R and D2R MSNs, both of which are targeted by HPC axons (34, 79, 80), might play dichotomous roles in appetitive and consummatory behaviors (see also refs (34, 81)). Future studies should aim to resolve the behavioral roles of specific NAc cell types targeted by HPC neurons.

In our analysis of dHPC^→NAc^ coding properties, we uncovered more neurons individually coding for two or three aspects of space, velocity, and lick activity specifically in identified projection neurons. Such conjunctive coding, or mixed selectivity, has been previously found in hippocampal neurons to combine coding of space with direction, velocity, appetitive behaviors, behavioral tasks, future paths (“splitter cells”), context, or objects (9, 12, 13, 55, 66, 82). These conjunctions are hypothesized to combine “where” and “what” information to disambiguate different experiences occurring in the same location, thus building a “scaffold” for episodic memories that allows unambiguous later retrieval (14, 83, 84). Relatedly, theoretical and experimental studies demonstrated that such high-dimensional coding facilitates action selection by putative downstream linear decoders tasked with action selection and generating behavior (61, 62, 85), and was found to scale with task demands and performance (55, 62, 85–87). Indeed, our data show that conjunctive coding improved reward zone detection by a linear decoder, similar to a recent study in retrosplenial cortex that demonstrated enhanced decoding near a reward zone by conjunctive coding neurons (87). In line with previous findings of increased conjunctive coding in prefrontal cortex → NAc projection neurons (88) and elevated proportions of splitter cells in dSub → NAc neurons (20), we thus propose that dHPC routes strongly conjunctive task-relevant information to facilitate NAc action selection.

The NAc has been described as a key node transforming motivational information from the limbic system into motor behaviors (25), but the information flow of specific projection neurons has remained elusive. Our data demonstrate that dHPC routes enhanced spatial information to the NAc that is conjunctively enriched by further spatial and non-spatial task-relevant features. We show that this conjunctive code improves linear decoding to guide downstream action selection, and that dHPC can drive the execution of appetitive motor behaviors via the basal ganglia, as early studies suggested (89). Thus, our findings identify a direct role for the hippocampus in the generation of motor behaviors and build an important bridge in our understanding of how sensory and mnemonic processes guide behavioral action.

## Materials and Methods

### Animals

Experiments were performed in adult male and female mice. C57Bl/6 (*n* = 16) and Thy1-GCaMP6s (*n* = 6) mice (GP4.3; The Jackson Laboratory, Bar Harbor, USA) were bred under specific pathogen-free conditions. Heterozygous mice were group-housed with 12 hours reversed dark light cycle at 21°C and ad libitum food/water access until mice had recovered from surgery. Experiments were performed during the dark phase. All experiments were performed according to the Directive of the European Communities Parliament and Council on the protection of animals used for scientific purposes (2010/63/EU) and were approved by the animal care committee of North Rhine-Westphalia, Germany.

### Viral vectors

To co-express the static red fluorophore mCherry in Thy1-GcaMP6s animals, we injected retrograde (rg) serotype (90) adeno-associated virus (AAV) under control of the polyglycokinase (pgk) promoter (AAVrg-pgk-Cre, Catalog #24593-AAVrg, titer: 1 × 10^13^ viral genomes (vg) ml^-1^, Addgene, Watertown, USA) in the NAc to enter axons and induce Cre expression in NAc-projecting neurons. Then, we used double-floxed inverse open reading frame (DIO) Cre-dependent mCherry under the human synapsin promoter in dHPC (AAV5-hSyn-DIO-mCherry, titer: 1.1 × 10^13^ vg/ml, Catalog #50459-AAV5, Addgene, Watertown, USA). For optogenetic experiments, we used unfloxed ChR2 (91) or control EYFP in excitatory neurons under the CaMKII promoter in the dHPC of C57Bl/6 mice (AAV2-CaMKII-hChR2(H134R)-EYFP-WPRE, titer: 4 × 10^12^ transducing units (TU), UNC #AV4381E, Vector Core at the University of North Carolina, Chapel Hill, USA; rAAV2-CaMKII-EYFP, titer: 4.3 × 10^12^ TU, UNC #AV6650, Vector Core at the University of North Carolina, Chapel Hill, USA).

### Stereotactic virus injections

For stereotactic injection of AAVs, mice were anesthetized with an intraperitoneal (i.p.) injection of a mixture of ketamine (0.13 mg/g) and xylazine (0.01 mg/g). Mice were head-fixed using a head holder (MA-6N, Narishige, Tokyo, Japan), placed into a motorized stereotactic frame (LuigsNeumann, Ratingen, Germany) and warmed by a self-regulating heat pad (Fine Science Tools, Heidelberg, Germany). After skin incision (5 mm) and removal of the periosteum, placement of the injection was determined in relation to bregma. A 0.5 mm wide hole was drilled through the skull (Ideal micro drill, World Precision Instruments, Berlin, Germany). Stereotactic coordinates were taken from Franklin and Paxinos, 2008 (The Mouse Brain in Stereotaxic Coordinates, Third Edition, Academic Press). To induce retrograde labelling of NAc-projecting neurons in dHPC, 2 × 500 nl of AAVrg-pgk-Cre were injected into the ipsilateral (right) nucleus accumbens (−1.3 mm anterior-posterior, -1.0 mm lateral, 5.0 and 4.3 mm ventral, relative to Bregma) at 100 nl/min, using a UltraMicroPump, 34G cannula and Hamilton syringe (World Precision Instruments, Berlin, Germany). To label projection neurons in red, 200 nl of AAV5-hSyn-DIO-mCherry were injected into the ipsilateral dorsal hippocampus (3.38 mm anterior-posterior, -2.5 mm lateral, 1.8 mm ventral, relative to Bregma, at a 10° angle). For optogenetic experiments, 200 nl of either AAV2-CaMKII-hChR2(H134R)-EYFP-WPRE or rAAV2-CaMKII-EYFP were injected each bilaterally into dorsal hippocampus (3.38 mm anterior-posterior, ± 2.5 mm lateral, 1.8 mm ventral, relative to Bregma, at a 10° angle). After surgery, buprenorphine (0.05 mg/kg) was administered thrice daily for 3 consecutive days. Implant surgery followed two weeks after AAV injection.

### Cranial window surgery

For awake hippocampal Ca2+ imaging, a window was surgically implanted over the right dorsal hippocampus, one week after virus injections. The hippocampal window was assembled from a 1.7 mm long stainless-steel cannula (3 mm outer diameter) and a round cover slip (3 mm diameter). The coverslip was glued to the end of the cannula using UV curable adhesive (NOA81, Thorlabs, Dachau/Munich, Germany). Mice were anesthetized and prepared for surgery as described above. The skin over the parietal skull and the periosteum were removed and wound edges sealed with Vetbond tissue adhesive (3M Animal Care Products, St Paul, USA). Skull surface was roughened by briefly applying gel etchant phosphoric acid (37.5 %; Kerr Dental, Scafati, Italy), carefully washing the skull surface and applying two-component dental adhesive (OptiBond FL, Kerr Dental, Scafati, Italy). A circular piece of the skull (3 mm in diameter) centered over the injection was carefully cut out using a sharp drill (Ideal micro drill, World Precision Instruments, Berlin, Germany). The dura was removed with forceps and mild vacuum suction was used to slowly remove the cortex within the craniotomy. Blood and aspirated tissue were washed out using a constant flow of artificial cerebrospinal fluid (ACSF). Intracranial aspiration was continued until the external capsule was exposed. The external capsule remained intact. The hippocampal window was manually inserted. During continuous perfusion with ACSF, the hippocampal window was lowered until sealing onto the external capsule. The hippocampal window was fixated, and the craniotomy sealed using UV curable dental cement (Gradia Direct Flo, GC Corporation, Tokyo, Japan). An angular metal bar (Luigs & Neumann, Ratingen, Germany) for head fixation was placed paramedian on the skull. After surgery, buprenorphine (0.05 mg/kg) was administered thrice daily for 3 days.

### Behavioral task

3-4 days after recovery from surgery, mice were provided with spinning wheels in their cages and were placed under a reverse light/dark cycle. One week after surgery, food restriction and two-week habituation schedules were initiated. Mice were food-restricted by providing about 80 % of their measured daily food pellets every 24 h. In the course of this, mice lost about 10-20 % of their original weight before the start of training. Habituation consisted of progressive exposure of mice to manual handling by the experimenter, obtaining milk rewards through a metal cannula, gentle manual head fixation, the treadmill apparatus, and, finally, head-fixed running on an unmarked treadmill belt with random rewards provided through a metal cannula lick spout after licking on it. The self-propelled treadmill (Luigs & Neumann) consisted of three rotating cylinders covered by a 7 cm wide and 360 cm long textile belt (Luigs & Neumann) including six differently textured zones: horizontal and vertical glue stripes, glue dots, Velcro dots, vertical tape stripes and upright nylon spikes. The reward zone for imaging experiments was 30 cm long and was placed between the end of the horizontal glue stripes and the beginning of the vertical tape stripes. The position of the mouse was recorded via an optical sensor (Luigs & Neumann) measuring the rotation of the treadmill cylinder underneath the mouse. Lick signals were measured by an analog piezo sensor; one full belt rotation was measured by an optical infrared sensor. All signals were collected at 10 kHz by an I/O board (USB-6212 BNC, National Instruments, Austin, USA) and recorded using custom-written Python software. After two weeks of habituation, mice were placed daily for 15 minutes on a cued (see above) treadmill belt on which they needed to lick on the metal spout in this hidden reward zone to obtain a liquid reward in the form of condensed milk. Milk was released by a miniature peristaltic pump (RP-Q1-S-P45A-DC3V, Takasago Fluidic Systems, Nagoya, Japan) that was triggered by custom-written Python software via an I/O board (USB-6212 BNC, National Instruments, Austin, USA). After five days of training, calcium activity was recorded while mice performed the learned task.

### Infrared camera behavioral tracking

Headfixed mouse behavior was continuously monitored by simultaneously using two monochrome CCD cameras (Basler acA 780-75gm) positioned at approximately 15 cm from the mouse. To capture face dynamics, we used a high-resolution zoom lens (50 mm FL, Thorlabs MVL50TM23); for body dynamics, we used a wide-angle lens (12 mm FL, Edmund Optics #33-303). Infrared illumination was provided via two 850 nm LED arrays (Thorlabs LIU850A), and cameras were outfitted with 850/40 nm bandpass filters (Thorlabs FB850-40). Both cameras’ positions were aligned for each mouse before the start of recordings. Camera images were acquired at 25 or 75 Hz with 782 × 582 pixels using pylon Camera Software Suite (Basler), each frame triggered by TTL pulses from the recording software. Files were saved in compressed MP4 format before further processing.

### Two-photon calcium imaging

Two-photon imaging was performed using an upright LaVision BioTec (Bielefeld, Germany) TrimScope II resonant scanning microscope, equipped with a Ti:sapphire excitation laser (Chameleon Ultra II, Coherent, Santa Clara, USA) operated at 920 nm for GCaMP6s fluorescence excitation, a second 1045 nm fixed-wavelength laser (Spectra Physics HighQ-2-IR, Newport Corp., Irvine, USA) for mCherry fluorescence excitation, and a 16x objective (N16XLWD-PF, Nikon, Düsseldorf, Germany). GCaMP6s fluorescence emission was isolated using a band-pass filter (525/40, Semrock, Rochester, USA) and detected using a GaAsP PMT (H7422-40, Hamamatsu, Herrsching am Ammersee, Germany). mCherry fluorescence emission was isolated using a band-pass filter (590/40, Semrock, Rochester, USA) and detected using a GaAsP PMT (H7422-40, Hamamatsu, Herrsching am Ammersee, Germany). Both lasers were aligned such that they excited the same focal plane. Imspector software (LaVision BioTec) was used for microscope control and image acquisition. Image series (2 channels, 1024 × 1024 pixels or 512 × 512 pixels, 350– 850 μm square field of view) were acquired at 15.2 Hz or 30.5 Hz, respectively. Frame capture signals were recorded by an I/O board (USB-6212 BNC, National Instruments, Austin, USA) that allowed subsequent data synchronization.

### Optogenetic stimulation

For optogenetic experiments, directly following virus injections, the skin over the parietal skull and the periosteum were removed and wound edges sealed with Vetbond tissue adhesive (3M Animal Care Products, St Paul, USA). Skull surface was roughened by briefly applying gel etchant phosphoric acid (37.5 %; Kerr Dental, Scafati, Italy), carefully washing the skull surface and applying two-component dental adhesive (OptiBond FL, Kerr Dental, Scafati, Italy). Two small 0.5 mm wide holes were drilled through the skull (Ideal micro drill, World Precision Instruments, Berlin, Germany) and a two-ferrules fiber-optic cannula (TFC_200/230-0.37_5.5mm_TS2.0_FLT, Doric Lenses, Quebec, Canada) was implanted bilaterally on top of NAc (coordinates: +1.3 mm anterior-posterior, +/-1 mm lateral, -4.9 mm ventral from brain surface, relative to bregma). The craniotomy was then sealed using UV curable dental cement (Gradia Direct Flo, GC Corporation, Tokyo, Japan). An angular metal bar (Luigs & Neumann, Ratingen, Germany) for head fixation was placed paramedian on the skull. After surgery, buprenorphine (0.05 mg/kg) was administered thrice daily for 3 days. After one week of recovery, the food restriction and a two-week habituation schedules were started. After successful habituation, head-fixed mice received light stimulation via a fiber-coupled 473 nm diode laser (LuxX 473-80, Omicron-Laserage) through a branching fiberoptic patchcord (BFP(2)_200/220/900-0.37_2m_FCM*-2xZF1.25; Doric Lenses, Quebec, Canada), at 5 mW light intensity, measured at the fiber output. Custom-written Python scripts were used to send a trigger signal to an analog isolated pulse generator (Model 2100, A-M Systems, Sequim, USA) once animals entered a hidden optogenetic stimulation zone. The pulse generator produced 5 ms long pulses at 20 Hz for a maximum duration of 10 seconds or until the mouse passed beyond the stimulation zone.

### Behavioral analysis

Analog signals for position, licking, reward pump, full rotation of belt, and digital signals from infrared camera triggers, two-photon scanning triggers, and optogenetic stimulation triggers were collected at 10 kHz and saved as TDMS files. Data processing and analysis were performed in Python. Position signals in cm were reconstructed using the optical sensor detecting a full rotation of the treadmill belt. Velocity was calculated using a Kalman filter applied to the position signal. Behavioral data were then down-sampled to either match the sampling rate of camera tracking (25/75 Hz) for training data or to match the sampling rate of two-photon imaging (15/30 Hz), by using each time window’s arithmetic mean (velocity), median (position, lap number, camera/scanner trigger), maximum (reward pump, optogenetic trigger), or standard deviation (licking). Discrete lick events were detected using Scipy’s *find_peaks* function with a minimum temporal distance of 0.33 s and a dynamic minimum height threshold that was individually determined by inspecting the synchronized infrared camera video. Lick bouts were classified as lick events that were <2 seconds apart from one another. Appetitive lick onsets were defined as lick bout onsets that were preceded by at least 3 seconds absence of lick events. Consummatory licking onset was defined as the first lick event between 0.5 seconds up to 5 seconds after reward pump trigger. Rewarded laps were defined as laps in which the reward pump was triggered by the animal’s licking. Relative licking refers to the cumulative analog lick signal per position bin or reward zone. Successful laps were defined as laps in which mice showed at least 50 % of relative licking in reward and anticipation zones and received a reward. High success trials were defined as trials consisting of at least 50 % of successful laps.

### Calcium signal processing

Two-photon imaging data was processed using custom-written software in Python, largely based on CaImAn (v1.6.2; (44)). Green and red channel 16-bit TIFF stack files (512 × 512 or 1024 × 1024 pixels times 9,000 or 18,000 frames) were first resampled to 1 px/μm before motion-correcting the red static channel using NormCorre piecewise rigid (parameters: *max_shifts* = 40, *num_frames_split* = 2000, *overlaps* = 46, *splits_els* = 4, *strides* = 255). The motion-corrected red channel image was then averaged over t and used for later identification of mCherry-positive components. Motion correction vectors were then applied on the dynamic green channel before using constrained non-negative matrix factorization (CNMF) for cell segmentation. Resulting traces were detrended and deconvolved before filtering spatiotemporal components using quality criteria followed by identity- and behavior-blind manual curation based on visual inspection of spatial and temporal footprints and the quality of deconvolution. To identify mCherry-positive components, resulting spatial footprints were overlaid over the red channel average and a dynamic threshold applied to visually match the optimal signal discovery.

### Place field analysis

Continuous belt positions were binned into 45 bins of 8 cm length. For calcium signals, de-convolved events were used to avoid differential effects of GCaMP6s calcium signal tails at different animal velocities across space. Only time points with a velocity >2 cm/s were considered for spatial tuning analysis. Spatial information (SI) was calculated as follows (47):

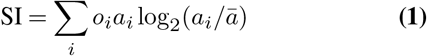

where *i* denotes the *i*-th spatial bin, *o*_*i*_ is the animal’s occupancy at spatial bin *i, a*_*i*_ is the mean of deconvolved events at spatial bin *i*, and *ā* is the overall mean calcium activity. Place cells were defined as cells whose SI was higher than the 95th percentile of 1000x randomly position-shuffled SI values (see ref (65)). Each shuffle consisted of randomly circularly shifting the activity time course by at least 500 frames, segmenting the activity in n blocks that were randomly permuted. n was automatically chosen for each session to match approximately twice the number of laps run to account for running differences between animals. The resulting shuffled trace was then used to calculate SI as above. Spatial information rate in bits per second was calculated by multiplying SI with the average firing rate during times of movement (>2 cm/s). Firing rate was defined as the number of deconvolved calcium events >0 per second. Sparsity was calculated according to ref (48), using the same denotation as above:

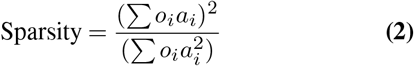

Place field reliability was calculated as the fraction of laps in which the maximum spatially binned deconvolved calcium activity occurred within the cell’s place field. i’.) in-out place field activity was defined as the average deconvolved calcium event activity within a cell’s place field subtracted by that activity outside the place field. Place fields were obtained by first averaging deconvolved calcium event activity at velocities >2 cm/s per 45 spatial bins, replicating the resulting trace by a factor of two to account for circularity of the belt, and applying a Savitzky-Golay filter (using Scipy’s *savgol* function with a window of 5 frames at the second order polynomial) to account for skewed place field activity. The resulting filtered spatial calcium activity was then searched for peaks using Scipy’s *find_peaks* function (*width*: 1 frame; *prominence*: 1.5 standard deviations; *relative height*: 0.8), with the most prominent peak used as primary place field and left/right interpolated intersection points as place field beginning/end. Place fields that would stretch into the replicated positions were translated back to the original 45 spatial bins. Place field boundary ratio was calculated as segmenting position bins into texture transition or middle zones, depending on their proximity to a texture transition, and calculating the ratio of place field beginnings/ends in a transition zone over those in a middle zone. To compare this ratio with a shuffled distribution, all place fields underwent 1000 iterations of randomly circularly shifting both beginnings and ends simultaneously and saving the resulting ratio both cell populations after each iteration. Place field centers were defined as the center of mass (COM) of significantly spatially modulated neurons (see above). For this, calcium activity at velocities >2 cm/s per 45 spatial bins was transformed to polar coordinates with q as position bin and r as average deconvolved calcium event activity at that position (see also ref (65)). The two-dimensional centroid was calculated, and the resulting angle transformed back to belt position to yield the place field center. “Place fields near reward zone” refers to place cells with COMs in either the anticipation zone (starting 30 cm before reward zone) or reward zone.

### Decoding of reward and anticipation zones

For decoding of reward and anticipation zones, all non-zero deconvolved calcium event activity was normalized into quantiles. Position and calcium activity were only considered from time points with velocity >2 cm/s. Data were downsampled to 1 Hz by accumulating calcium activity and averaging position data. Positions in the 30 cm reward zone or the preceding 30 cm anticipation zone were binarized (1 inside zone, 0 outside zone). Models were trained using calcium data of either (non-)projecting or (non-)conjunctive neurons and 100 randomly sample size-matched neurons of the respectively larger population, ultimately using the average *R*^2^ of the random sampling procedure. Neurons with fewer than 10 time bins of non-zero calcium activity in train/test datasets were excluded from training/testing of the model. A linear support vector machine (SVM) classifier (Sklearn’s *linear_model*.*SGDClassifier* function; *loss*: hinge, *penalty*: L2) was cross-validated by training on even laps and testing on odd laps, and vice versa, using the average of the resulting *R*^2^. For models comparing (non-)projection neurons only imaging sessions with >10 projection neurons were included.

### Velocity and lick coding

Velocity modulation was assessed by linear regression of each neuron’s average deconvolved calcium event activity against 1 cm/s velocity bins from 2 up to 30 cm/s using Scipy’s stats.linregress function (see also ref (56)). Cells were considered to be significantly velocity modulated at P_adjusted_ < 0.05 after correcting for false discovery rate by using Benjamini/Hochberg correction. Significantly velocity modulated cells with positive slopes were termed “speed-excited” and those with negative slopes “speed-inhibited”. To assess population-level appetitive/consummatory lick modulation, average deconvolved calcium activity before and after each lick event (see above for definition) were compared within the respectively described time windows. For single-cell level appetitive lick modulation analysis, each neuron’s average pre/post deconvolved calcium activity was compared across events using Wilcoxon’s non-parametric paired test. Cells were classified as “licking-excited” if they were significantly modulated and showed a positive difference, or “licking-inhibited” if they showed a significant negative difference.

### Generalized linear model

To determine cellular coding of space, velocity, and licking, we used a generalized linear model to predict each neuron’s calcium activity. We normalized calcium activity by dividing each neuron’s deconvolved calcium events by their signal-to-noise ratio and the resulting standard deviation. Normalized calcium activity was then smoothed using a Gaussian window of 0.5 seconds (*SD* = 2) and temporally downsampled by averaging to 3 Hz. Finally, downsampled non-zero calcium data were normalized into quintiles. The feature matrix consisted of normalized position, velocity, and lick data. Position data were binned into 45 equally sized spatial bins (8 cm each) and median-averaged to 3 Hz, used as factors. Velocity data were normalized by dividing data by its mean and averaging to 3 Hz. Lick bouts were calculated as described above. Appetitive lick bouts were defined as all lick bouts occurring up to 5 seconds before and at least 5 seconds after reward dispensation. Appetitive lick bouts were downsampled by median-averaging to 3 Hz. Both downsampled velocity and downsampled licking (appetitive lick bouts) were replicated with time shifts spanning 3 seconds before to 3 seconds after original timing at 3 Hz to account for anticipatory or delayed calcium activity. Generalized linear Poisson models were trained, validated and tested using Python’s H2O library (*H2OGeneralizedLinearEstimator*; *lambda*: 0) with a train/validation/test ratio of 0.8/0.1/0.1. The resulting *R*^2^ score based on the test dataset was saved and compared to that of randomly shuffled feature models. For each predictor group (position, velocity, licking), 100 shuffled models were generated by circularly shifting the respective group’s data randomly between 200 and 700 time points, saving each model’s *R*^2^ score. Significant predictors for each neuron’s calcium activity were defined as those whose random shuffling procedure resulted in reductions in *R*^2^ for more than 95 % of cases. This means that at least 96 of the 100 models with one randomly shuffled predictor group had to have decreased *R*^2^ values compared to the full model, in order for this predictor to be considered significant. Conjunctive-coding neurons were defined as those with at least two significant predictor groups.

### Infrared camera recordings analysis

Markerless pose estimation (Deep Graph Pose (92) and DeepLabCut (93)) was used to detect facial and body movements for both types of videos. For this, a deep neural network was trained to automatically discriminate 15 markers for videos of the body (paw, tail and head segments) and 13 markers for videos of the face (6 for pupil, eye, nose, mouth, etc). The network was trained on a large variety of lighting conditions and angles until it reached satisfactory performance. Mouth facial regions of interest (ROIs) were automatically segmented using video-averaged marker points of nose tip, the eye’s tear duct and the mouth as stable landmarks. Mouth motion energy was calculated as the ROI’s average pixel-by-pixel intensity differences from *t*_−1_ to *t*_0_, and *z*-scoring the resulting activity.

### Statistical analysis

Statistical analysis was performed using Python’s pingouin library. Statistical tests are indicated in the figure legends and text. To evaluate statistical significance, data from Figures 1D and 3F was subjected to one-or two-way ANOVAs. Data in Figures 2D, 4D, 4H, 5H-I, 7B, 7D, 7F, 8B-C, and 8E was subjected to chi-square tests. Data in Figures 2E-H was subjected to Welch’s *t*-tests. Data in Figures 3C-D was subjected to bootstrapping analysis (see above) and chi-square tests. Data in Figure 3G and 8G was subjected to Wilcoxon’s test for nonparametric data. Data in Figure 5C was subjected to a repeated-measures ANOVA followed by post-hoc *t*-tests with Bonferroni correction. Data in Figures 6I and 6K was subjected Student’s paired t-test. For all analyses data are presented as mean ± SEM, and the threshold for significance was at *p* < 0.05.

## ACKNOWLEDGEMENTS

The authors would like to thank Janelle Pakan, Hugo Spiers, Magdalena Sauvage, Marlene Bartos, and Thomas Mrsic-Flögel as well as the rest of the Cellular Neuroscience Department for their insightful comments and discussions while preparing this manuscript. In addition, the authors would like to thank Falko Fuhrmann, Pavol Bauer, and Rüdiger Geis for their help with the treadmill setup, cranial surgery, and two-photon imaging. The resources for all experiments were provided by the German Center for Neurodegenerative Diseases (DZNE) in Bonn and the Leibniz Institute for Neurobiology (LIN) in Magdeburg. The authors received funding from the German Research Foundation (DFG) SFB grant 1089 (P.M., S.R.) and ERC Consolidator grant Sub-D-Code (O.B., S.R.).

## AUTHOR CONTRIBUTIONS

O.B. and S.R. conceived and designed experiments. O.B. performed and analyzed imaging experiments. O.B. and P.M performed optogenetic experiments, O.B. analyzed these. O.B. wrote the manuscript with help from S.R. and P.M..

## CODE AND DATA AVAILABILITY

Code and data will be made available upon final publication.

## Supplementary videos

**Supplemental Video 1. *In vivo* dual-color two-photon calcium imaging of a mouse during goal-directed navigation**. Representative example of dual-color two-photon calcium imaging combined with goal-directed navigation and lick behavior. Infrared camera images show face (top left) and body (top right) tracking. Calcium activity is shown by overlaying denoised GCaMP6s activity (green) onto average mCherry signal (red; bottom left); traces represent denoised GCaMP6s activity of mCherry-negative (green) and mCherry-positive (red; NAc-projecting) hippocampal neurons (bottom right). Playback speed is 2*×* original speed.

**Supplemental Video 2. Optogenetic stimulation induces mouth movement**. Five representative optogenetic stimulation trials from one experimental mouse expressing ChR2 and one control mouse expressing EYFP only. A light fiber in NAc stimulated excitatory hippocampal projections by shining 473 nm laser light at times indicated by blue squares (top) and blue traces (bottom). Optogenetic stimulation repeatedly induces lick behavior and a decrease in velocity in ChR2-expressing animals but not those expressing EYFP. Playback speed equals original speed.

**Figure S1.**
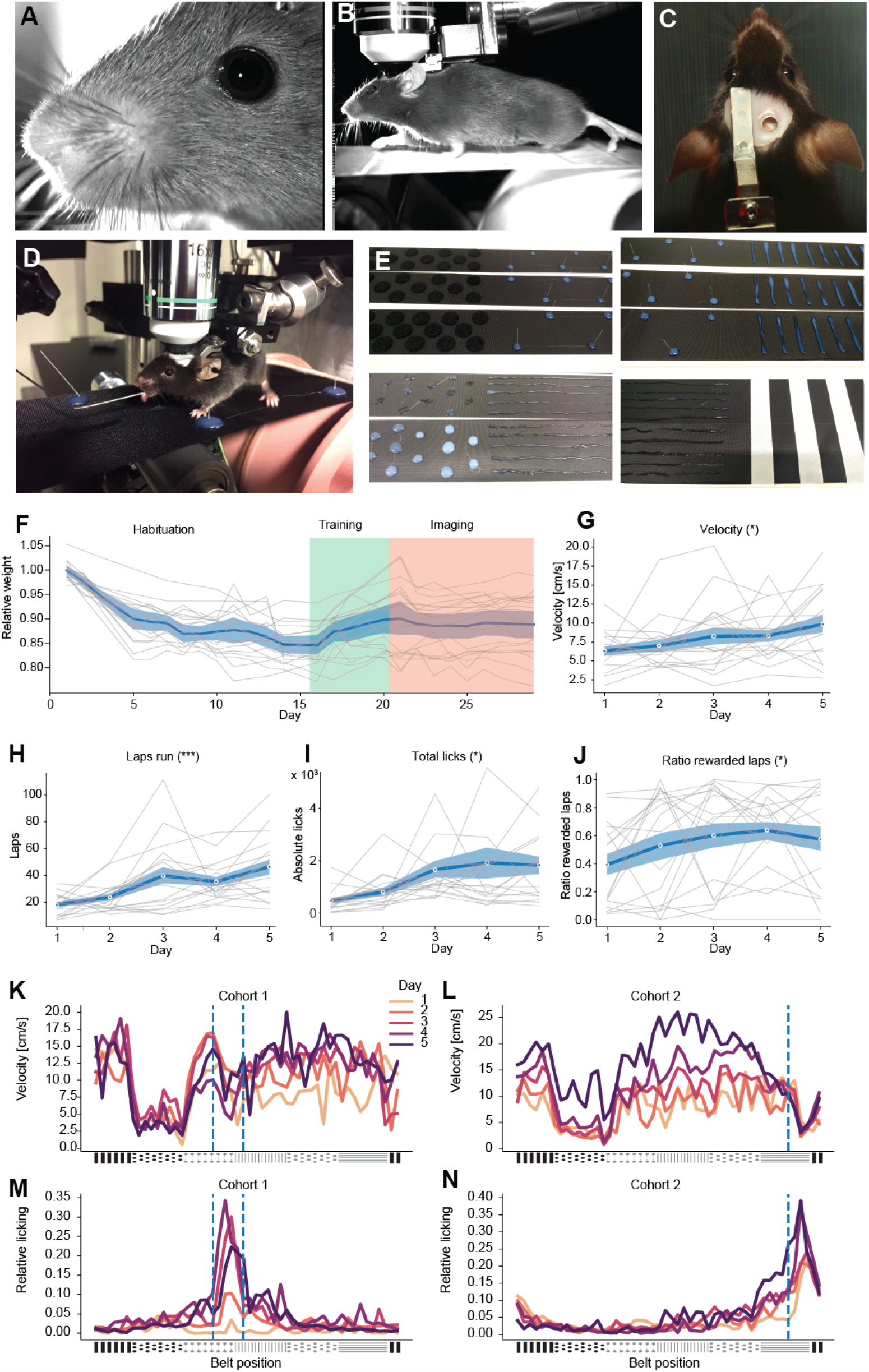
Spatial reward learning task. (A and B) Sample infrared images of face (A) and body (B) that were continuously captured at 25/75 Hz. (C) Image of one head-fixed experimental mouse from top, illustrating craniotomy and head-fixation. (D) Sample image of two-photon imaging of a head-fixed mouse on treadmill belt, licking at the lick spout. (E) Sample images of the six different belt texture zones, including Velcro dots, nylon spikes, vertical glue stripes, glue dots, horizontal glue stripes, white tape stripes (from left top to right bottom). (F) Weight changes after introduction of food restriction and during training and imaging. (G-J) Average velocity (G, *F* (4) = 2.631, *p* = 0.0430), number of laps run (H, *F* (4) = 8.771, *p* < 0.001, GG-corrected), number of licks (I, *F* (4) = 3.883, *p* = 0.0326, GG-corrected), and ratio of rewarded laps (J, *F* (4) = 3.331, *p* = 0.0157) increase over the course of five training days (all repeated-measures ANOVAs). Gray lines indicate data points of individual animals, blue shade represents SEM, blue line represents mean. (K-N) Average velocity (K and L) and average licking (M and N) changes across belt position and days of mice tested with reward zone in “center” of belt (K and M) and “end” of belt (L and N). Blue dashed line represents presence of reward zone. *n* = 9 (cohort 1), *n* = 9 (cohort 2). All data are presented as mean ± SEM. \**p* < 0.05, \*\*\**p* < 0.001.

**Figure S2.**
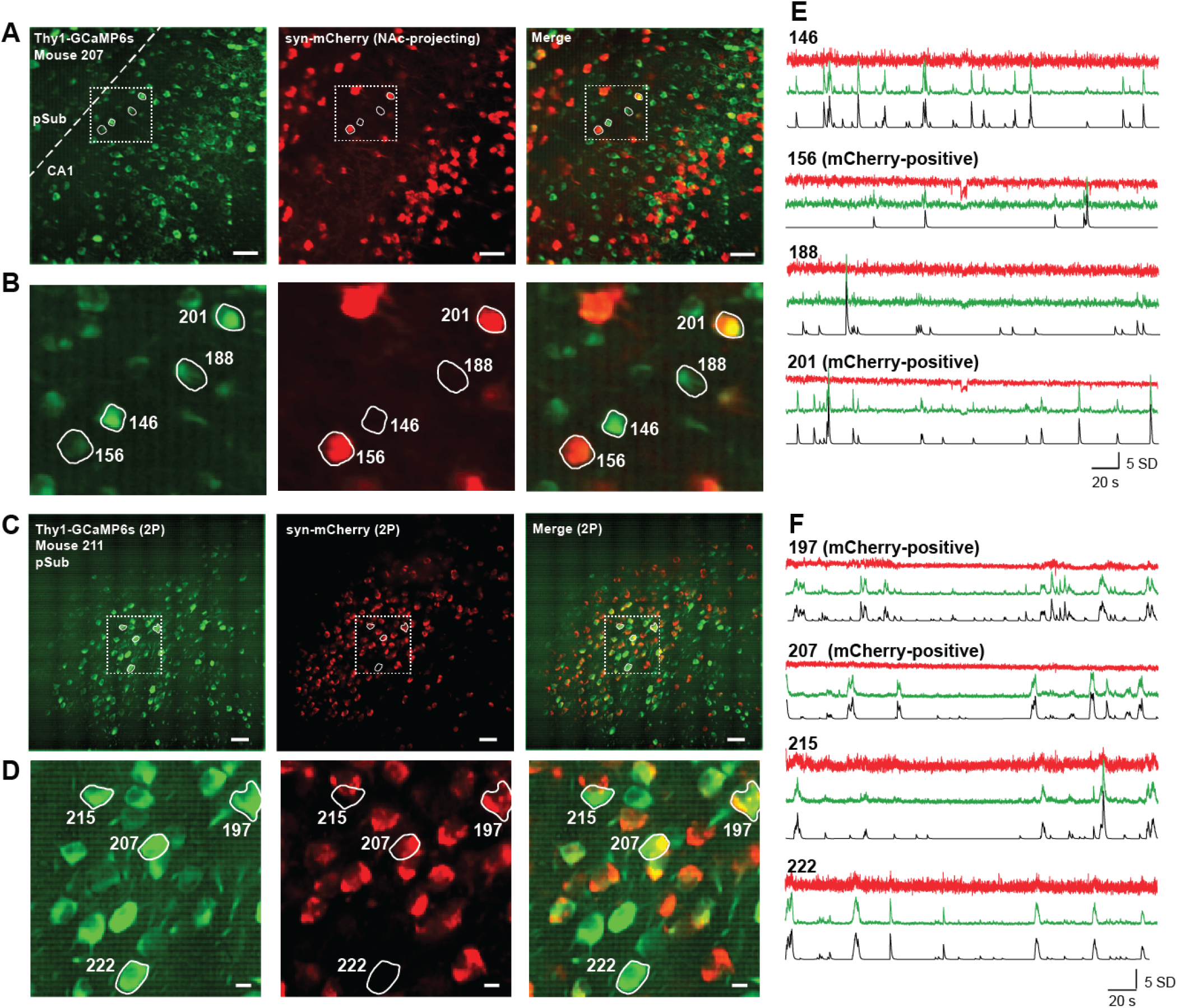
*In vivo* dual-color two-photon imaging of dynamic GCaMP and static mCherry signals. (A) Sample field of view (FOV) of one imaged region crossing putative CA1 and prosubiculum (pSub). Green channel shows GCaMP6s local correlations image, red channel shows average motion-corrected mCherry signal. (B) Detail of region in (A) clearly outlining spectrally separate channels. (C and D) Sample FOV of another region in putative prosubiculum (C) with detailed view (D). Scale bars represent 50 μm (A and C) or 10 μm (B and D). (E and F) Z-scored fluorescent temporal sample traces of regions of interest (ROIs) shown in (A-D). Red traces show raw fluorescence dynamics from imaging red channel, green traces show raw fluorescence dynamic from green channel, and black traces show denoised calcium signal.

**Figure S3.**
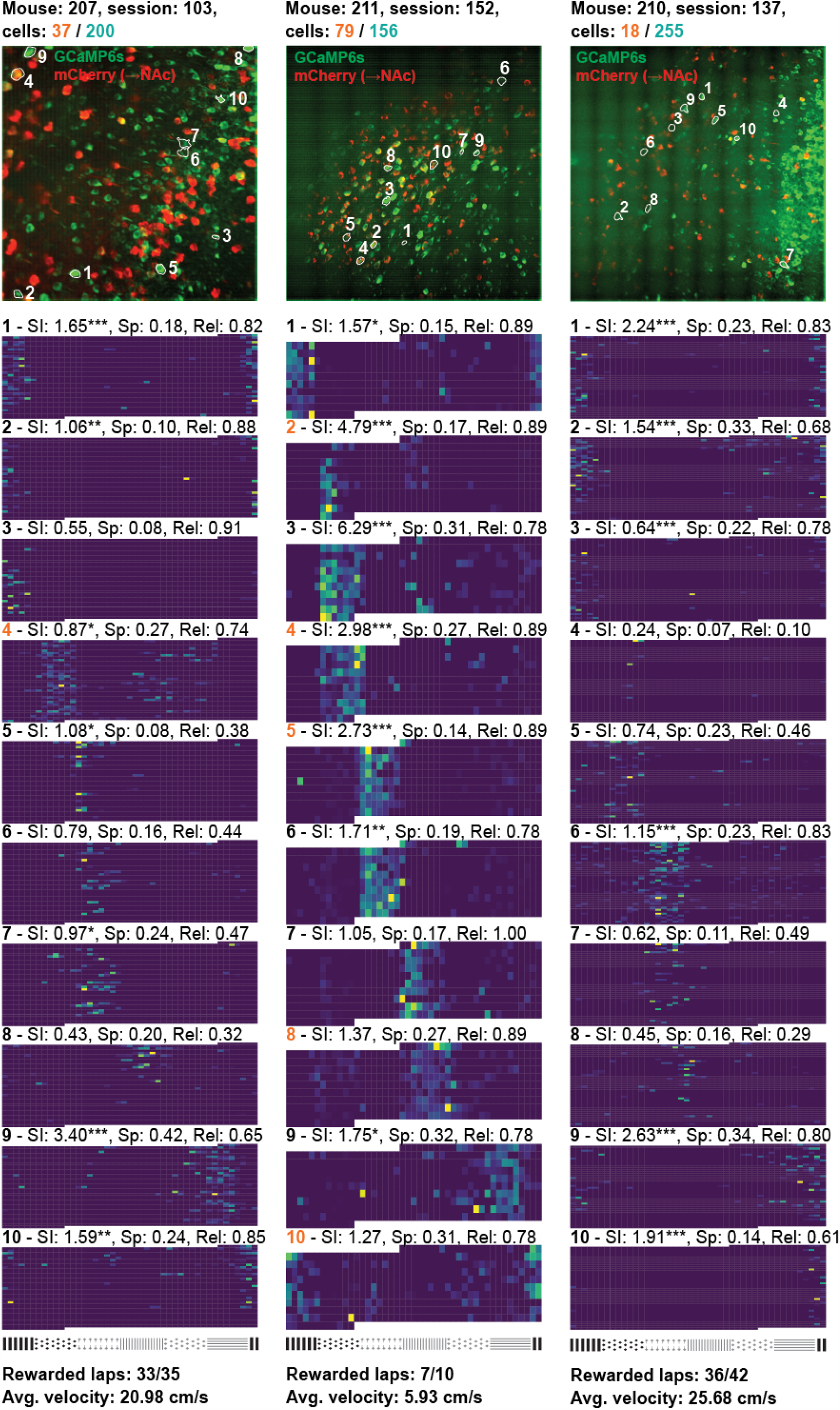
Spatial tuning of different neurons in sample FOVs. Three representative fields of view (FOVs) from three different mice are shown, with single-lap spatial calcium activity of each 10 representative neurons, some putatively NAc-projecting. FOV information is shown on top, including numbers of identified dHPC^−^ (green) and dHPC^→NAc^ (red) neurons. FOVs shown are composites of CaImAn maximum local correlation images (*caiman*.*summary_images* .*max_correlation_image*) of GCaMP channel (green) and the averaged motioncorrected red channel with NAc-projecting mCherry fluorescence. Contours of ten representative neurons from each FOV are indicated with white outlines and numbers that refer to spatially averaged calcium activity below. Each neuron’s normalized average calcium activity across 45 spatial bins per lap (y axis) is shown with key spatial information values above (SI: spatial information (47), Sp: Sparsity (48), Rel: reliability of each neuron’s per-lap maximum activity to occur within the place field). Orange numbers refer to NAc-projecting neurons.

**Figure S4.**
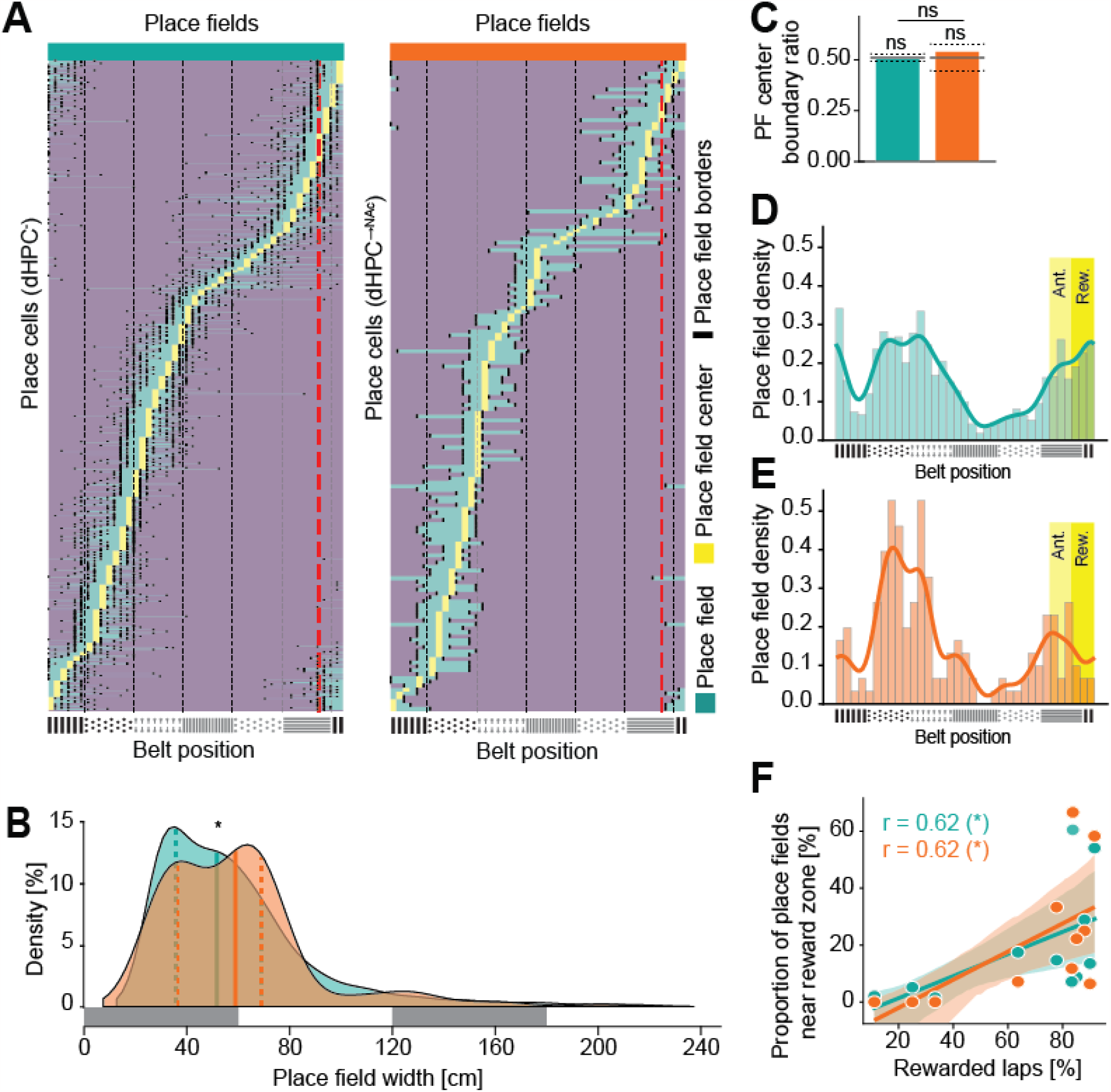
Place field distribution across the belt and behavioral performance. (A) Extent of place fields of dHPC^−^ (left) and dHPC^→NAc^ (right) place cells, sorted by each place cell’s center of mass (COM). Turquoise represents place field, yellow represents COM, black lines represent left and right borders of place fields. Red dashed line represents beginning of reward zone, black dashed lines represent belt texture borders. (B) Width of place fields is unevenly distributed between dHPC^−^ (green; quartiles 35.6 cm / 51.7 cm / 69.0 cm) and dHPC^→NAc^ (red; quartiles 36.5 cm / 58.9 cm / 69.0 cm) place cells (Kolmogorov-Smirnov test, D = 0.116, *p* = 0.0299). Gray bars represent belt texture zones (60 cm). (C) Place field centers are not biased with respect to texture boundaries (percentile < 95th, permutation test) and the ratio is not different between neuronal populations (χ^2^(1, 5372) = 1.646, *p* = 0.1995). (D and E) Place field density between reward and anticipation zones compared to the rest of the belt for dHPC ^−^ (D) and dHPC^→NAc^ (E) populations. (F) Reward and anticipation zone overrepresentation correlates with behavioral success (percentage of rewarded laps per session) for both dHPC ^−^ (green) and dHPC^→NAc^ (red) populations (Pearson correlation; *r* _dHPC–_ (15) = 0.622, *p* = 0.0107; 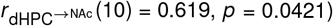. ns: not significant, \**p* < 0.05.

**Figure S5.**
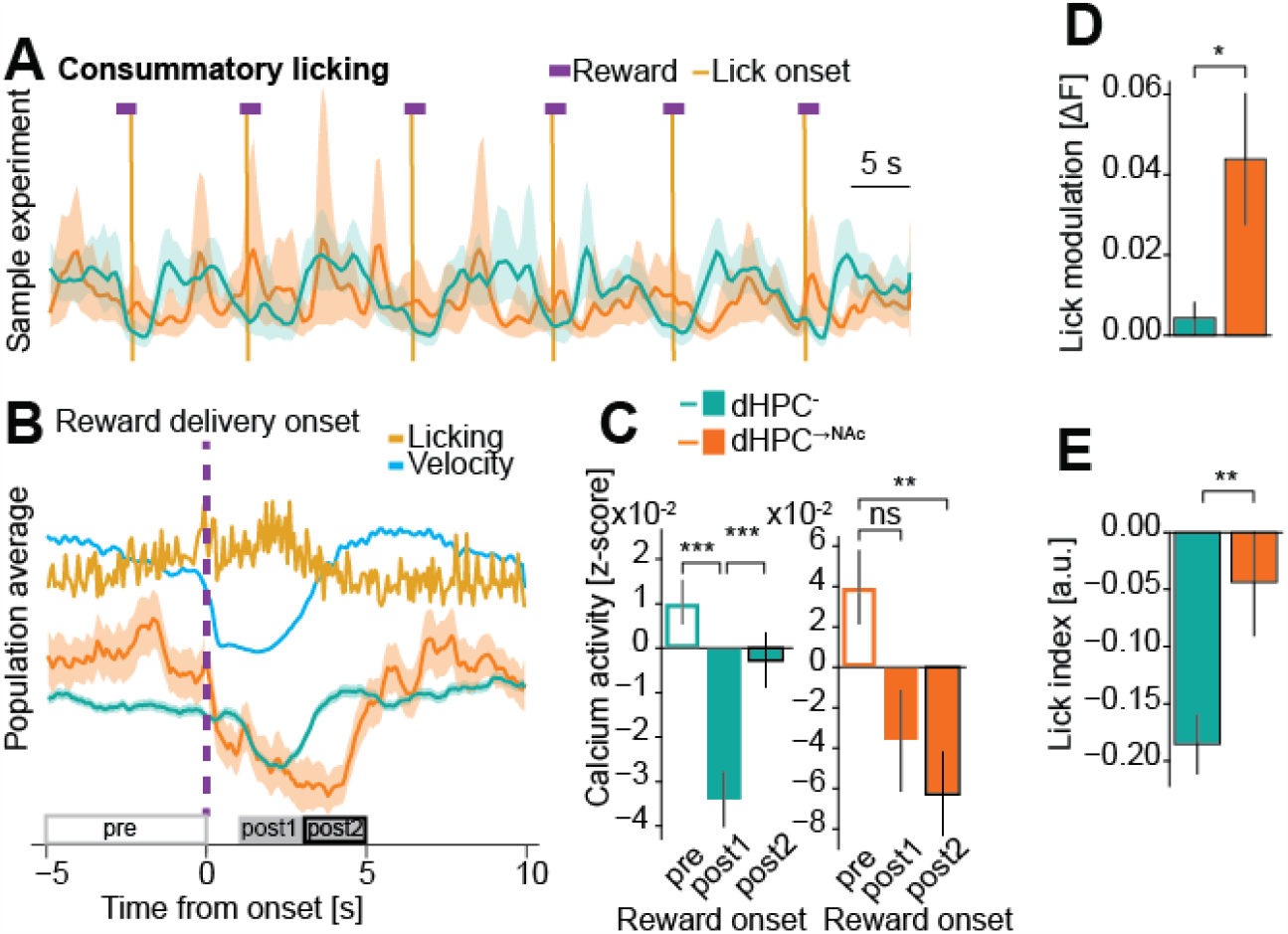
Lick modulation during consumption and general licking. (A-C) Consummatory licking leads to a depression in neural activity. (A) Sample experiment showing reward dispensation (purple bars) and consummatory licking onsets (golden vertical lines) and population calcium activity from dHPC^−^ (green) and dHPC^→NAc^ (red) neurons. Note the robust depression of neural activity during times of reward consumption. (B) Event-triggered average traces around reward delivery onset, including licking and speed as well as population average calcium activity of dHPC^−^ (green) and dHPC^→NAc^ (red) neurons. Gray rectangles indicate time windows for comparing calcium activity shown in (C). (C) Calcium activity is differentially modulated by reward onset (two-way mixed ANOVA; *F* _reward_timing_ (2, 10740) = 19.380, *p*(GG-corrected) < 0.001, *F* _projection_(1, 5370) = 0.798, *p* = 0.372, *F* _interaction_ (2, 10740) = 5.559, *p* = 0.0039). Post-hoc pairwise *t*-tests with Bonferroni correction were performed for each interaction and the results indicated with asterisks. (D) Lick modulation (average of each trial’s pre-post difference) is significantly increased in dHPC^→NAc^ neurons (Welch’s *t* (496.11) = 2.379, *p* = 0.0177). (E) Lick index (average calcium activity difference during licking) is significantly increased in dHPC^→NAc^ neurons (Welch’s *t* (454.19) = 2.729, *p* = 0.0066). All data are presented as mean ± SEM. \**p* < 0.05, \*\**p* < 0.01, \*\*\**p* < 0.001.

**Figure S6.**
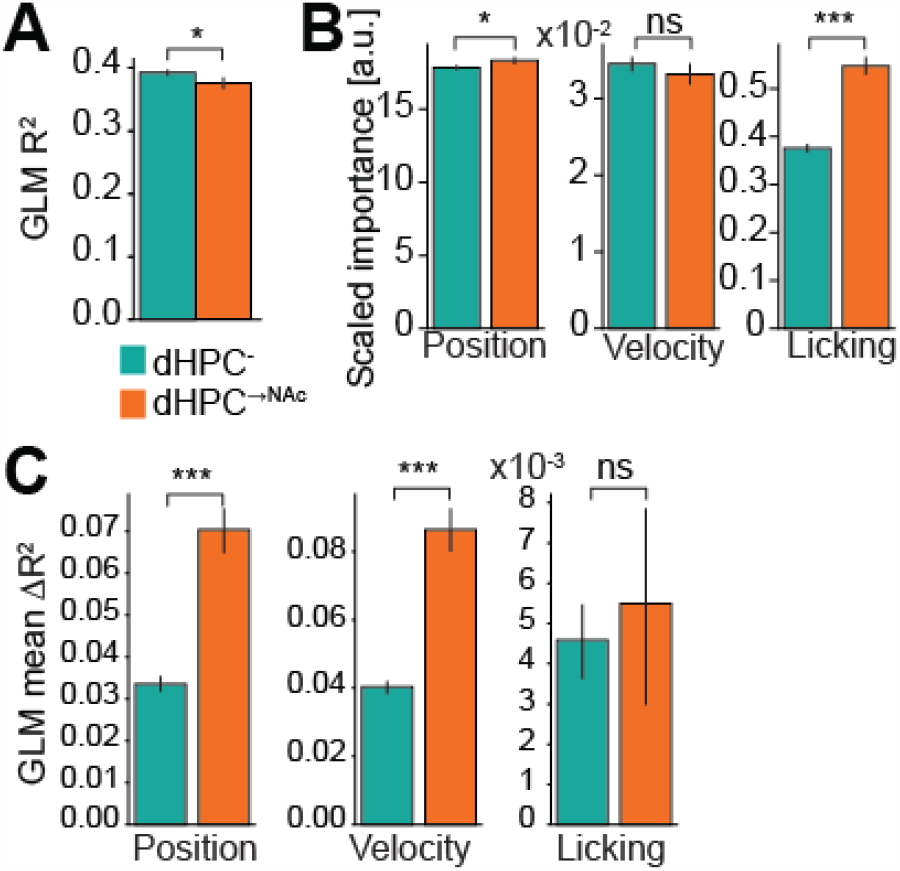
Generalized linear model reveals coding differences for dHPC^→NAc^ neurons. (A) GLMs’ explained variance *R*^2^ is significantly lower for dHPC^→NAc^ neurons (Welch’s *t* (615.15) = 2.147, *p* = 0.032). (B) Scaled feature importance (standardized GLM coefficients retrieved by H2O’s varimp() method) is higher in dHPC^→NAc^ neurons for position (Welch’s *t* (2508.09) = 2.162, *p* = 0.031) and licking (Welch’s *t* (1879.75) = 9.328, *p* < 0.001), but not for velocity (Welch’s *t* (2609.15) = 0.881, *p* = 0.378). (C) Mean *R*^2^ differences between full and feature-shuffled models are significantly greater in dHPC^→NAc^ neurons for position (Welch’s *t* (523.23) = 5.984, *p* < 0.001), velocity (Welch’s *t* (497.69) = 7.704, *p* < 0.001), but not for licking (Welch’s *t* (552.63) = 0.3626, *p* = 0.717). All data are presented as mean ± SEM. ns: not significant, \**p* < 0.05, \*\*\**p* < 0.001.

